# Viral but not bacterial community succession is characterized by extreme turnover shortly after rewetting dry soils

**DOI:** 10.1101/2023.02.12.528215

**Authors:** Christian Santos-Medellín, Steven J. Blazewicz, Jennifer Pett-Ridge, Joanne B. Emerson

## Abstract

As central members of soil trophic networks, viruses have the potential to drive substantial microbial mortality and nutrient turnover. Pinpointing viral contributions to terrestrial ecosystem processes remains a challenge, as temporal dynamics are difficult to unravel in the spatially and physicochemically heterogeneous soil environment. In Mediterranean grasslands, the first rainfall after seasonal drought provides an ecosystem reset, triggering microbial activity during a tractable window for capturing short-term dynamics. Here, we simulated precipitation in microcosms from four distinct, dry grassland soils and generated 144 viromes and 84 metagenomes to characterize viral, prokaryotic, and relic DNA dynamics over 10 days. Vastly different viral communities in each soil followed remarkably similar successional trajectories. Wet-up triggered a significant increase in viral abundance and richness, followed by extensive compositional turnover. While temporal turnover in prokaryotic communities was much less pronounced, differences in the relative abundances of Actinobacteria (enriched in dry soils) and Proteobacteria (enriched in wetted soils) matched those of their predicted phages, indicating viral predation of dominant bacterial taxa. Rewetting also rapidly depleted relic DNA, which subsequently re-accumulated, indicating substantial new microbial mortality in the days after wet-up, particularly of the taxa putatively under phage predation. Production of abundant, diverse viral particles via microbial host cell lysis appears to be a conserved feature of the early response to soil rewetting, and results suggest the potential for ‘Cull-the-Winner’ dynamics, whereby viruses infect and cull but do not decimate dominant host populations.

## Introduction

Viruses drive microbial predator-prey dynamics and contribute to community turnover and biogeochemical cycling on global scales^1, 2^. In the oceans, viruses kill up to 40% of their microbial hosts each day, with consequences for food web dynamics, carbon and nutrient cycling, and the maintenance of biodiversity^1–4^. With up to 10^10^ viral particles per gram of soil, viruses are assumed to be influential in terrestrial ecosystems too^5–10^, but viral impacts on soil ecology and biogeochemistry are just beginning to be explored. Although methodological barriers for soil viral community analyses have largely been overcome^10–13^, the recent advent of large-scale viral size-fraction metagenomics (viromics) in soil has heralded new challenges associated with extreme soil viral diversity^12, 14–18^.

Given viral dependence on hosts for replication, soil viral dynamics are inherently linked to host abundance and activity. Yet, despite co-enrichments of microbial taxa and their viruses under selective environmental conditions^19, 20^, beta-diversity patterns of soil viral and prokaryotic communities often show significant differences. In particular, spatial structuring has recently emerged as a major factor influencing soil virosphere assembly^15–18^, with viral community distance-decay relationships often eclipsing those of bacteria and archaea^15, 16^. Whether these differences in viral and microbial spatial turnover also apply over temporal scales is unknown. Further, while significant differences in soil viral community composition and/or abundance have been observed over months^14, 15, 21–23^, these relatively long-term dynamics are difficult to decouple from changing environmental conditions and could also be confounded by spatial structuring^8, 16^. Evaluating temporal trajectories in the context of spatial heterogeneity therefore requires high spatiotemporal resolution, ideally under controlled environmental conditions.

In the Mediterranean climate zone - where long, dry summers are followed by cool, wet winters - the first annual rainfalls trigger a predictable, near-immediate efflux pulse of carbon dioxide and a burst of microbial activity and carbon mineralization^24–29^. This annual ecosystem reset has lent itself to tractable, controlled microcosm experiments, which have consistently shown that soil rewetting reshapes the prokaryotic composition of soil microbiomes, with extensive turnover of active taxa during the first few days following soil rewetting^30–34^. In support of a concomitant viral response to wet-up, substantial increases in viral diversity were observed in biocrust soils upon rewetting^35^. Conversely, a decrease in viral richness and a steady increase in viral biomass were observed in grassland microcosms following wet-up, and subsequent successional dynamics were evident but partially obscured by spatial structuring in the field from which the soils were derived^18^. Teasing apart robust viral successional patterns post-wet-up thus requires both biological replication across soils and technical replication across independent microcosms.

Viromics, arguably the best method for studying soil DNA viral communities^14, 15, 36^, can also offer a window into ‘relic’ DNA^37^ dynamics, convenient for exploring microbial mortality patterns. On average, 50% of the extractable DNA in soil has been shown to be ‘relic’ DNA, or free, extracellular DNA^37–39^. While most studies that have considered relic DNA have focused on understanding and/or reducing its impact on microbial community measurements in support of targeting the active microbiome^37–39^, relic DNA itself may offer a window into microbial mortality, including in response to viral predation^16, 40^. The composition of relic DNA can be inferred by first removing it through laboratory treatments and then comparing resultant sequencing libraries to those from total DNA^37, 39^. However, the post-0.2 µm (‘viromic’) size fraction offers the potential to measure relic DNA more directly^16^. Briefly, purified viral particle (virion) fractions are typically treated with DNase to remove relic DNA prior to virion lysis, but skipping this DNase treatment leaves a virome that retains both viral DNA and relic DNA that co-precipitated with virions^16, 36^. By accessing this relic DNA pool, DNase-untreated viromes may yield insights into microbial mortality patterns not apparent in total DNA^16^, providing a more holistic view of community dynamics.

Here we leveraged DNase-treated viromes, DNase-untreated (hereafter, untreated) viromes, and total metagenomes from four distinct grassland soils to interrogate viral and microbial successional and mortality patterns in 144 microcosms following wet-up. A more thorough understanding of the coupled responses of prokaryotes and their viruses to environmental cues, in the context of the spatiotemporally, physicochemically, and biologically complex soil environment, is essential for better predicting soil microbiome dynamics in a changing world.

## Results

### Dataset overview and key features, including compositional differences by both omic analysis method and soil type

Near-surface soils (0-15 cm) were collected at the end of the 2020 dry season (soil moisture content 1% or lower) from four grassland habitats in the Mediterranean climate zone in northern California (**Figure 1a, Supplementary Figure 1a-b**). Soils were sourced from three sites (Jepson, Hopland, and McLaughlin), and Jepson soils were collected from two topographical features: mima mounds (small, dome-like structures) and their adjacent swales (shallow depressions that fill with water during winter as vernal pools) (**Figure 1a**). For each of the four soil types, dry soils were added to highly replicated microcosms and rewetted to ∼25% soil moisture content with sterile water (**Supplementary Figure 1c**). Microcosms were destructively sampled at seven time points over ten days: T0 (dry) and T1-T6 (24, 48, 72, 120, 168, and 240 hours post-wet-up, respectively), and a control set of unwetted soil microcosms was also sampled after 10 days **(Figure 1b**). To characterize the short-term temporal dynamics of the soil viral community response to rewetting, as well as compare temporal patterns in bacterial and viral communities and relic DNA following wet-up, we generated 144 viromes (84 DNase-treated and 60 untreated) and 84 total metagenomes (**Figure 1c**).

**Figure 1.**
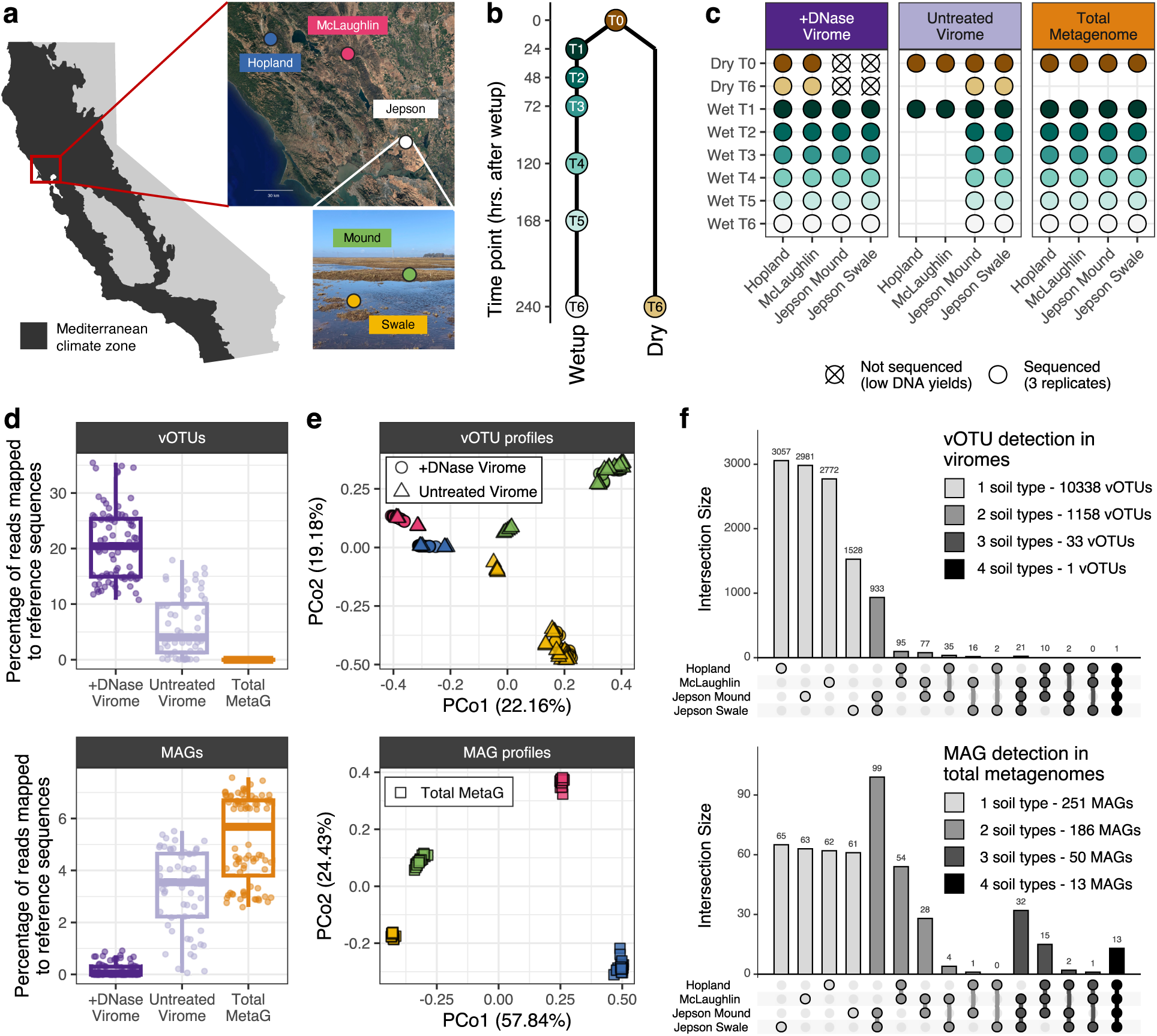
Dataset overview. **(a)** Sources of the four grassland soils used in this study: map shows the distribution of the Mediterranean climate zone in California, USA; zoomed area shows the locations of the three grassland sites; photograph shows the two topographical features sampled in the Jepson site (photo from the rainy season to highlight differences between mounds and swales; soils were collected when dry). **(b)** Time points in this study: top point (T0) corresponds to the pre-wet-up dry soils at the beginning of the experiment, left points (T1-T6) correspond to the six time points after wet-up; right point (T6) corresponds to the control dry soils collected at the end of the experiment. **(c)** Sets of triplicate total metagenomes and viromes (DNase-treated and untreated): filled circles correspond to three sequenced replicates; crossed-out circles indicate triplicate samples with DNA yields too low for library construction. **(d)** Percentage of quality-filtered reads mapped to the sets of dereplicated vOTUs (upper facet) and MAGs (lower facet). Boxes show the median and interquartile range. **(e)** Unconstrained analyses of principal coordinates performed on vOTU Bray-Curtis dissimilarities calculated across viromes (upper facet) and MAG Bray-Curtis dissimilarities calculated across total metagenomes (lower facet). Colors indicate soil type (as in panel a). **(f)** Upset plots of detection patterns across soil types for vOTUs in viromes (upper facet) and MAGs in metagenomes (lower facet); circles indicate soil types for each detection pattern, and intersection size is the number of vOTUs or MAGs exhibiting a particular detection pattern.

We identified a total of 11,533 viral operational taxonomic units (vOTUs, ≥10 kbp genome fragments, ≥95% average nucleotide identity^41^) and 513 metagenome-assembled genomes (MAGs, ≥50% completeness, ≤10% contamination). Based on the proportion of reads that mapped to vOTUs versus MAGs, the three omic sample types reflected a continuum of viral content that was highest in DNase-treated viromes, intermediate in untreated viromes, and lowest in total metagenomes; that pattern was mirrored in reverse for prokaryotic content (Figure 1d, **Supplementary Figure 2**). Both viral and prokaryotic communities differed most significantly among the four soils (**Figure 1e**), with 90% of vOTUs and 50% of MAGs exclusively found in microcosms from one of the four soil types (**Figure 1f**). These minimal overlaps in community composition confirm that we captured substantially different grassland soil habitats (biological replicates) for temporal analyses.

### Wet-up triggered diverse viral particle production across soils, with the magnitude of viral richness increase tied to precipitation history

Despite vastly different composition across soils, viral communities followed remarkably similar responses to wet-up. For all four soil types, rewetting resulted in a significant increase in viral richness within 24 hours, which remained comparatively high for the rest of the experiment (**Figure 2a**). In contrast, prokaryotic (MAG) richness was relatively unchanged upon wet-up and throughout the 10-day experiment (**Figure 2a**). Further, the increase in viral diversity triggered by rewetting was accompanied by an increase in viral particle (virion) abundance in all four soils, measured by proxy as DNase-treated viromic DNA yields (**Figure 2b**).

**Figure 2.**
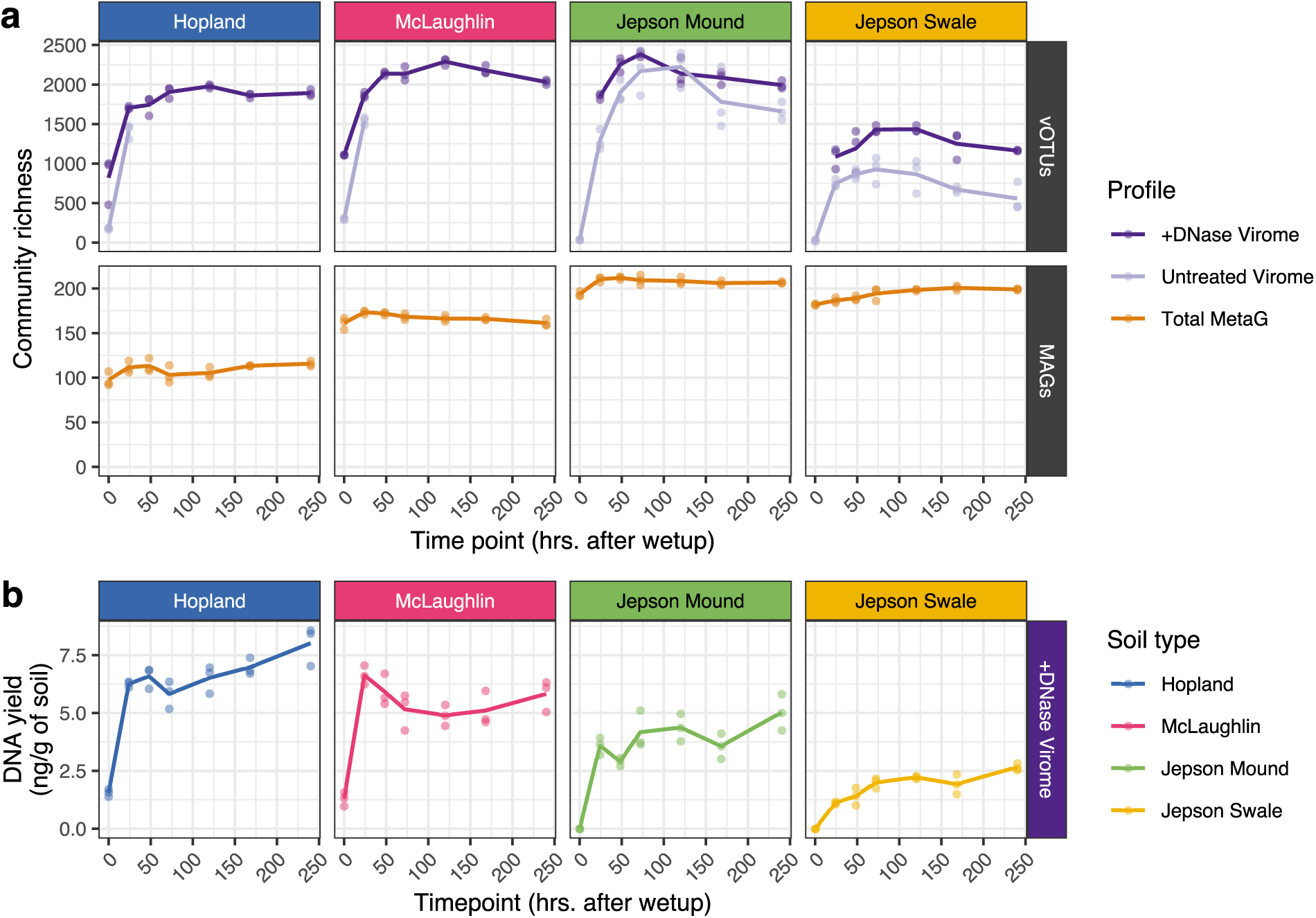
Richness and virion abundance over the time series. **(a)** Community richness measured as the number of vOTUs (top facet) or MAGs (bottom facet) recovered from soil microcosms: points correspond to individual viromes or total metagenomes, and trend lines track the mean richness across replicates at each collection time point. **(b)** Temporal trends in DNase-treated viromic DNA yields (a proxy for virion abundance). Points correspond to individual microcosms, and trend lines track the mean yields across replicates. Yields were below detection limits for pre-wet-up samples from Jepson soils (represented as zero in the graph).

The magnitude of increased viral richness after wet-up differed by soil type, with field precipitation patterns prior to sample collection potentially explaining the differences. Both Jepson soil types displayed a >30-fold increase in detected vOTUs 24 hours after wet-up, compared to only a ∼2- fold increase for Hopland and McLaughlin soils (**Figure 2a**). In dry soils, both Hopland and McLaughlin had substantially greater viral richness than Jepson (**Figure 2a**), as well as more viral particles measured by viromic DNA yields (**Figure 2b**). While no major rainfall was registered at Jepson over the 208 days before our samples were collected, it rained >5 mm 74 days before sample collection at both Hopland and McLaughlin (**Supplementary Figure 1b**). The more prolonged dry period at Jepson could have depleted virions from the preceding rainy season, while the higher viral richness in Hopland and McLaughlin dry soils could have resulted from remains of a viral bloom triggered by summer rainfall. To further explore links between dry soil viral richness and precipitation history, we consulted dry and rewetted soil viromic data from Hopland in the preceding two dry seasons (2019 and 2018). Consistent with increased virion depletion after a longer dry period, Hopland soils that experienced a 150-day dry period in 2019 had much lower viral richness (consistent with dry Jepson soils here) than the 2020 Hopland soils here collected only 74 days after rainfall (Supplementary Figure 3a-b), while soils from 2018 collected 96 days after a ≥5 mm rain event (Supplementary Figure 3b) had high viral richness^18^. Precipitation history thus seems to have had measurable impacts on dry soil viral diversity and the magnitude change in viral richness in response to wet-up.

Given that virion (free viral particle) abundance was below detection limits in two of the dry soils and that 81% of vOTUs observed after rewetting were not detected in dry soil, virions may not be the primary dry season reservoir of soil viral populations. To evaluate whether prophages (i.e., viral genomes integrated in host microbial genomes via lysogeny) were the likely reservoir, we classified vOTUs with integrases and/or excisionases in their annotations as putative temperate phages (capable of lysogeny). Across all four soils, temperate phages accounted for only 9.7% of vOTUs, and their aggregated relative abundances remained below 13.6% (**Supplementary Figure 4**), suggesting that most vOTUs produced upon wet-up were not capable of lysogeny. Thus, neither lysogeny nor the presence of free viral particles (except perhaps at low abundance below detection limits) can readily explain the survival of a reservoir of highly diverse viral populations during the dry season.

### Post-wet-up viral communities exhibited significant temporal succession, which was less pronounced in prokaryotic communities

For each soil type, the initial spike in viral richness after wet-up was followed by compositional succession over the subsequent nine days (**Figure 3a**). As viral communities from the control dry soil microcosms at the end of the experiment (T6) were statistically indistinguishable from those at T0 (**Figure 3a, Supplementary Table 1)**, the observed successional patterns resulted from the water addition. In all cases, the most significant difference in viral community composition was between dry soils and all post-wet-up time points, consistent with the biggest shift in viral diversity observed within the first 24 hours of rewetting. After 24 hours, wetted soils displayed steady changes in viral beta-diversity that were reproduced across biological (different soil) and technical (triplicate microcosm) replicates and that were robust to differences in sample processing (DNase-treated and untreated viromes). In short, post-wet-up dynamics were reproducibly characterized by extensive turnover of soil viral communities.

**Figure 3.**
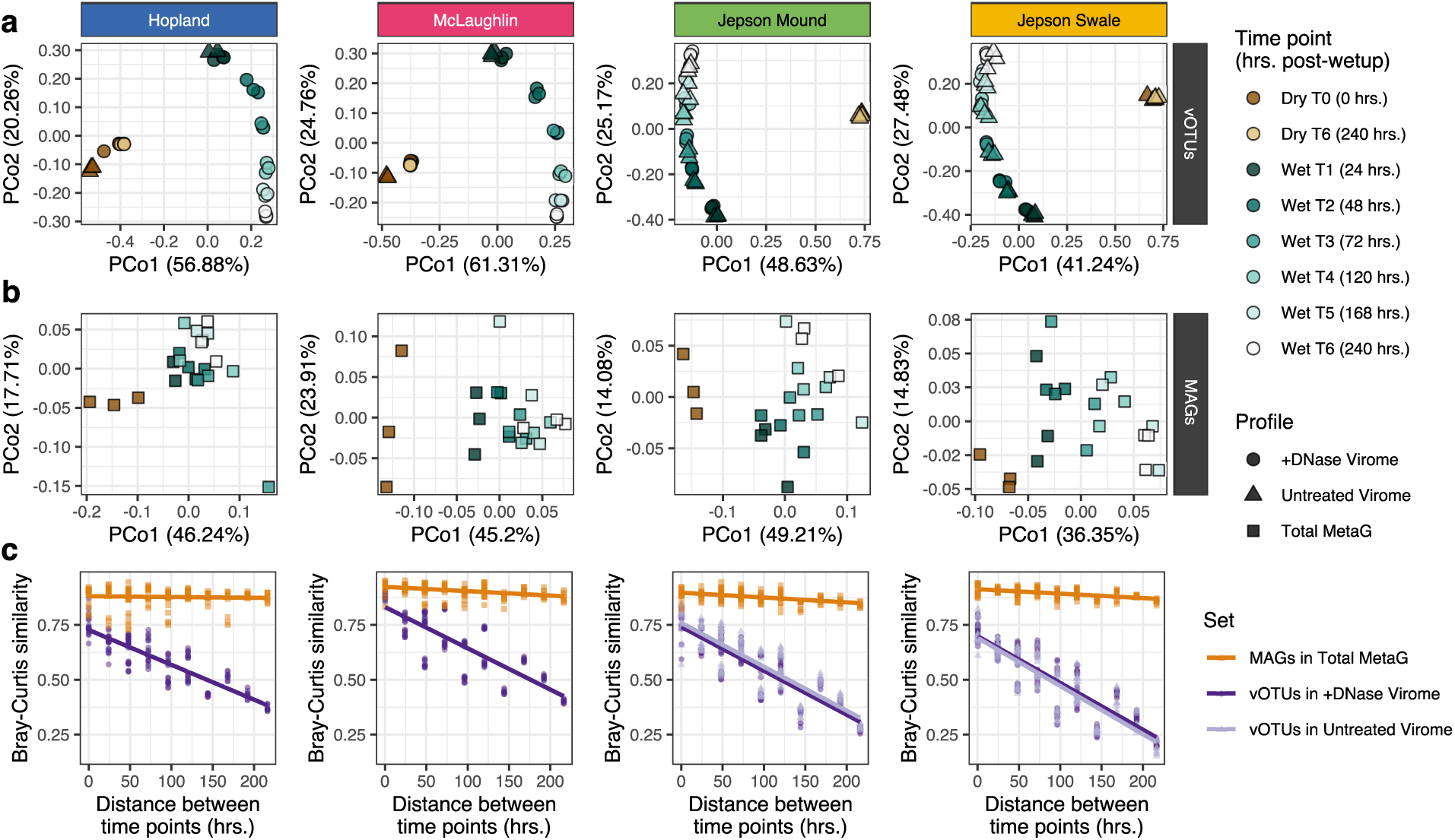
Successional patterns across the four soil types for viral compared to prokaryotic communities. **(a,b)** Unconstrained analyses of principal coordinates performed separately for each soil type on **(a)** vOTU Bray-Curtis dissimilarities from viromes and **(b)** MAG Bray-Curtis dissimilarities from total metagenomes. **(c)** Linear regressions between Bray-Curtis similarities and temporal distances in post-wet-up time points for viral (vOTUs in viromes) and prokaryotic (MAGs in total metagenomes) communities. Points represent pairs of samples, trend lines show the least squares linear regression model, and facets correspond to soil types. MetaG = metagenome.

Bacterial and archaeal communities were also affected by the rewetting event, but the magnitude of their response was not as pronounced as for the viruses. To study prokaryotic communities, we considered both MAGs and 16S rRNA gene amplicon profiles from total DNA. Briefly, results from the 16S rRNA gene analyses were similar to those from the MAGs (**Supplementary Figure 5, Supplementary Table 2**), so we focus here on the MAGs. While changes in prokaryotic (MAG) community composition between dry and wet soils were consistent across the four soil types, turnover across post-wet-up time points was not as significant as for the viral communities (**Figure 3b**). On average, 82% of MAGs were recovered in all six post-wet-up time points per soil, relative to only 35% of vOTUs (**Supplementary Figure 6a**). Differences in the degree of temporal turnover in prokaryotic and viral communities were exemplified by the overlap in populations between 24 hours and 10 days after wet-up (T1 and T6), which was 73% for MAGs and only 15% for vOTUs (**Supplementary Figure 6b**). Across all four soil types for viruses and only three for prokaryotes, Bray-Curtis community similarity was significantly negatively correlated with temporal distance (**Supplementary Table 2**). In all cases, the temporal distance-decay relationship was far more significant for viral communities, as reflected in much steeper slopes, with viral community temporal turnover being on average eight times higher (**Figure 3c, Supplementary Table 2**). The emerging picture is of much more significant spatial^15, 16^ and, here, temporal turnover of viruses compared to their prokaryotic hosts in soil.

### The viral response to rewetting included taxonomically conserved dynamics, reflecting predation of dominant bacterial taxa

The consistent pattern of extensive viral community turnover within each soil suggested the potential for a taxonomically and/or functionally conserved, trait-based viral response across soils. We identified 6,220 vOTUs whose abundances in DNase-treated viromes differed significantly across time points in at least one soil type and grouped them into three trait groups: dry-dominant (vOTUs at relatively high abundance in dry soil, decreasing upon wet-up), early responders (vOTUs peaking in abundance 24-48 hours post-wet-up), and late responders (vOTUs generally increasing from low to high abundance over the course of the experiment) (**Figure 4a**). Only 215 of these temporally dynamic vOTUs were detected in multiple soils, but of those, 93% were in the same trait group (**Supplementary Figure 7**), suggesting that ‘species-level’ viral responses were consistent across soils.

**Figure 4.**
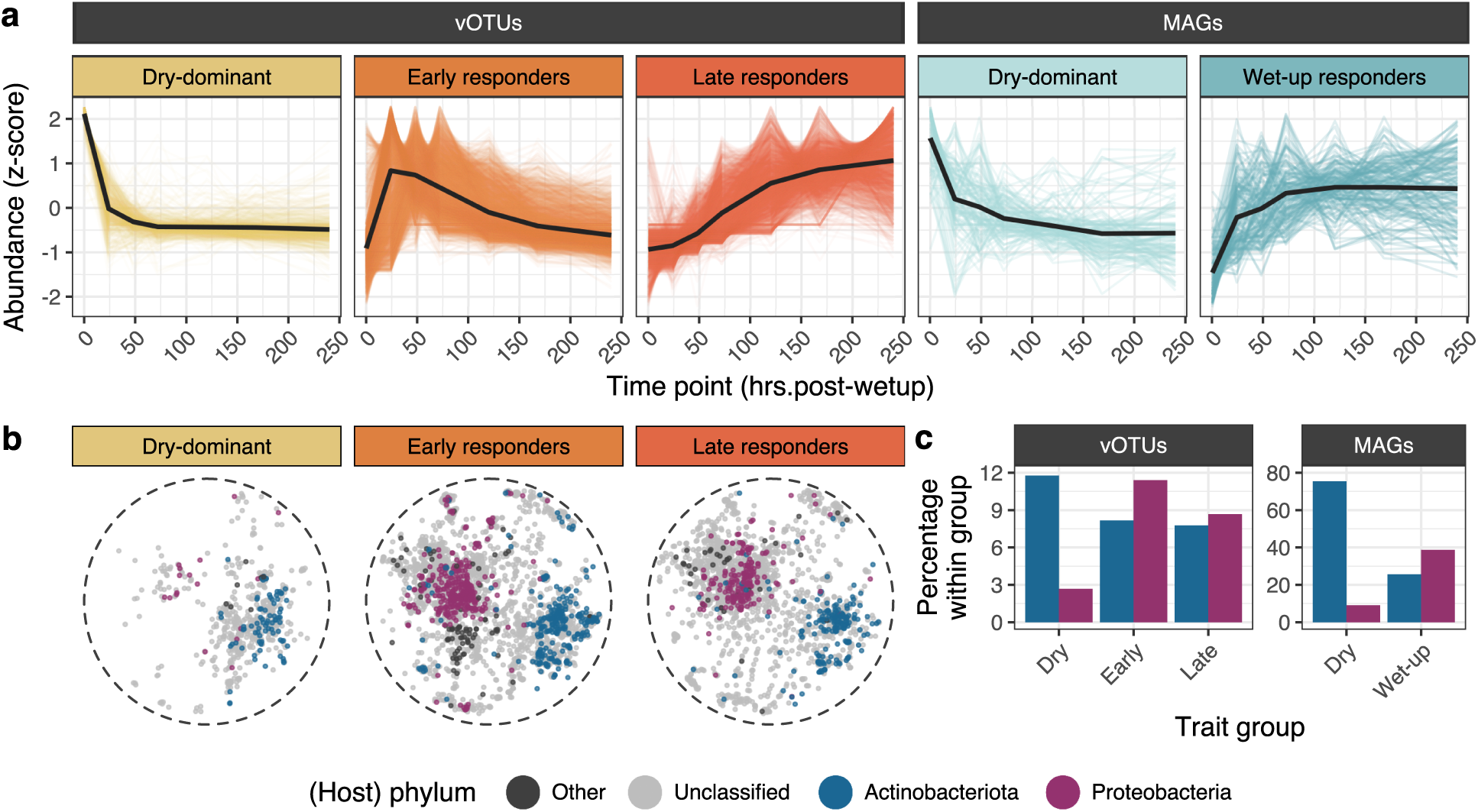
Temporal trait-based and taxonomic patterns in the viral and prokaryotic responses to wet-up across soils. **(a)** Temporal trends in the abundances of differentially abundant vOTUs in DNase-treated viromes and MAGs in total metagenomes from all four soils. Colored trend lines represent the z-transformed mean abundances of single vOTUs or MAGs across time points, bold black trend lines correspond to the mean abundances of the subset of vOTUs or MAGs included in each temporal trait group (facet). **(b)** Distribution of the subset of vOTUs in each temporal trait group within the predicted protein-sharing network of all 11,533 vOTUs from all four soils (Supplementary Figure 8b). Each node is one vOTU in a Fruchterman-Reingold network layout constructed from pairs of vOTUs with a significant overlap in their predicted protein contents. Color indicates predicted host taxonomy. **(c)** Percentage of vOTUs (left facet) and MAGs (right facet) classified as (or with hosts classified as) Actinobacteria and Proteobacteria in each temporal trait group. Full taxonomic trends can be found in **Supplementary Figures 8b-c** (vOTUs) **and 9** (MAGs).

We next investigated whether conserved viral responses to wet-up extended to higher taxonomic levels, potentially allowing for exploration of virus-host dynamics. Considering all 11,533 vOTUs (from all soils at all time points), we identified vOTUs with significant overlap in their predicted protein contents, reflecting genomic relatedness^16, 42^, using network analysis (**Supplementary Figure 8b**). To determine whether viral responses to wet-up were taxonomically conserved, we compared clustering patterns in the protein-sharing network for each trait group (**Figure 4b**). Indeed, hypergeometric tests revealed different regions of the network as significantly over- and underrepresented for each trait group, indicating that genomically related vOTUs tended to respond similarly to wet-up, regardless of soil type (**Supplementary Figure 8a**). Given that viral taxonomic networks are coarsely structured by host taxonomy^15, 16, 42, 43^, we next wondered whether the observed shifts in viral ‘taxa’ (network regions) between trait groups could be explained by shifts in the predominant host(s) preyed upon across the time series. Leveraging host taxonomic assignments via iPHoP^43^ (**Supplementary Figure 8b**), significant shifts in predicted hosts were observed across trait groups (**Supplementary Figure 8c**, **Figure 4c**). While putative actinobacteria-infecting phages were proportionately distributed within the early and late responder trait groups, they were significantly overrepresented in the set of dry-dominant vOTUs, suggesting the preferential infection of actinobacteria under dry (or drying) soil conditions prior to the start of the experiment. In contrast, putative Proteobacteria phages were significantly underrepresented in the dry-dominant trait group but became significantly overrepresented among early responders, hinting at increased infection and lysis of Proteobacteria hosts shortly after rewetting.

We next sought to determine whether these temporal trends in predicted hosts matched those of the putative hosts themselves, for example, reflecting viral predation of abundant prokaryotic taxa. As MAGs were detected more consistently across time points than vOTUs (**Supplementary Figure 6a**), they separated best into two temporal trait groups: dry-dominant (abundant in dry but generally not wet soil) and wet-up-responsive (depleted in dry soil and relatively abundant throughout most or all of the wet time points) (**Figure 4a**). Consistent with more putative actinophages and fewer putative proteobacteriophages observed in dry soil, significantly more Actinobacteria and significantly fewer Proteobacteria were in the dry-dominant compared to wet-up-responsive MAG groups (**Figure 4c, Supplementary Figure 9**). Similarly, the increase in the abundance of putative proteobacteriophages upon rewetting coincided with an overrepresentation of Proteobacteria MAGs in the wet-up-responsive group. Thus, the dynamic taxonomic composition of viral communities in response to wet-up broadly tracked with shifts in the relative abundances of host taxa, consistent with viral infection of dominant bacteria.

### Relic DNA as both fuel for microbial activity and a window into microbial mortality

We leveraged our untreated viromes, which included extracellular DNA^37^ due to the lack of DNase treatment^36^, to investigate relic DNA dynamics and microbial mortality. Yield comparisons between DNase-treated and untreated viromes revealed that DNase-digestible DNA comprised the majority of the post-0.22-µm size fraction (**Supplementary Figure 10**). For all four soil types, untreated viromic (relic) DNA yields were highest in dry soils and dropped precipitously within 24 hours of wet-up (**Figure 5a**), suggesting that relic DNA accumulated in dry soils and was degraded quickly after rewetting. For Jepson microcosms, which had untreated viromes from all time points, untreated viromic DNA yields remained well above DNase-treated viromic DNA yields throughout the experiment (**Supplementary Figure 10**), indicating substantial relic DNA at all time points. Relic DNA started to re-accumulate towards the end of the experiment (**Figure 5a**), suggesting new microbial mortality later in the time series. Overall, these results hint at both microbial mortality throughout the experiment and a potential role of accumulated relic DNA from the dry season as a nutrient-rich substrate and/or source of nucleic acids^39, 44, 45^ that contributed to the burst of microbial activity immediately after rewetting.

**Figure 5.**
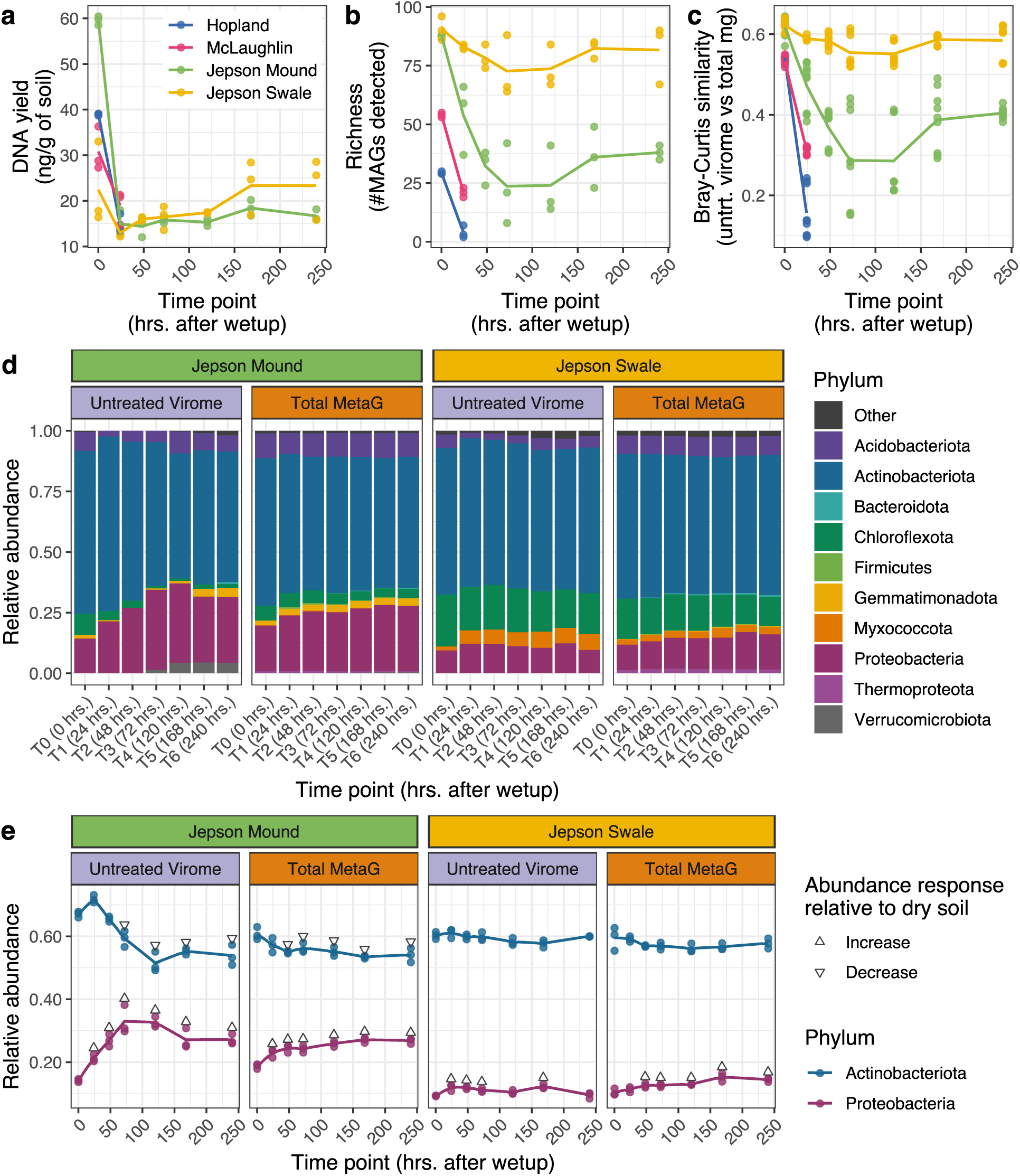
Relic DNA abundance and composition over time. **(a-b)** Temporal trends in **(a)** DNA yields and **(b)** the number of MAGs recovered from untreated viromes by soil type. For **(a)** and **(b)**, points correspond to individual samples, and trend lines represent the mean across replicates at each time point. **(c)** Prokaryotic compositional similarities (pairwise MAG community Bray-Curtis similarities) between untreated viromes (relic DNA, untrt. virome) and total metagenomes (total mg) over time. Points are sample pairs, and trend lines follow the mean Bray-Curtis similarity at each time point. **(d)** Taxonomic composition of MAGs in untreated viromes and total metagenomes from Jepson soils. Stacked bars represent the mean relative abundances of MAGs aggregated at the phylum level. **(e)** Aggregated relative abundances of Actinobacteria and Proteobacteria MAGs in untreated viromes and total metagenomes from Jepson soils. Points correspond to individual microcosms, and trend lines track the mean abundances across replicates. Triangles indicate a significant increase or decrease in abundance relative to dry soils (adjusted P-value < 0.05, Dunnett’s test). For panels **(b-e)**, we excluded MAGs detected in DNase-treated viromes, as they were likely derived from intact ultra-small cells present in the 0.22 µm fraction (not representative of relic DNA).

We next analyzed the prokaryotic composition of relic DNA to reveal which microbes were dead or dying over time. We focused on 249 MAGs detected in untreated viromes (**Supplementary Figure 2b**), excluding 20 MAGs that were also recovered from DNase-treated viromes (**Supplementary Figure 2b**), which could represent intact (non-relic) ultra-small bacteria (in particular, ultra-small Patescibacteria^46^ were enriched in our DNase-treated viromes, **Supplementary Figure 11**). Consistent with the observed high relic DNA abundance in dry soil and its rapid depletion upon wet-up, dry soils had among the highest numbers of MAGs recovered in relic DNA, and a significant decrease in relic DNA MAG richness was observed 24 hours after rewetting for all four soil types (**Figure 5b**). Where untreated viromes were available throughout the time series, MAG richness in relic DNA increased during the second half of the experiment (**Figure 5b**), suggesting that extracellular DNA in wet soil reflected new mortality of diverse microorganisms a few days after rewetting. As the observed viral community dynamics suggest extensive infection and lysis of microbial hosts throughout the experiment, viral lysis could be one mechanism that would explain continued microbial mortality over 10 days.

Whether virus-induced mortality tends to drive turnover within stable host populations or leads to more drastic decimation is an open question that warrants a comparison of the total (metagenomic) and dead (relic DNA) microbial fractions. We recognize that microbes captured in total metagenomes could exist anywhere along the continuum from actively growing to dormant to dead, so here we simply seek to compare broad compositional patterns for likely living and likely dead groups of microbes. Total metagenomes would share the same composition as relic DNA either if the same taxa were living and dead, or if the relic DNA pool were so substantial that it overwhelmed the signal from living biota. Here, relic and total community MAG profiles were most similar in dry soils (**Figure 5c**), consistent with a relic DNA pool so large that DNA from dead organisms dominated the total metagenomes. Relic and total MAG composition diverged immediately following wet-up for all four soil types, potentially reflecting high microbial replication (i.e., genomes from fresh biomass eclipsing the relic DNA signal in the total metagenomes) and low mortality in the first days after wet-up. For the Jepson soils with untreated viromes throughout the time series, relic and total MAG composition trended towards increasing similarity over the final four days of the experiment, suggesting that more of the same microbes were replicating and dying over time and/or that more of the relic DNA pool was contributing to the total DNA pool.

Despite some differences in their degrees of similarity over time, overall, the total and relic DNA pools were largely similar in composition, especially at coarse taxonomic resolution (**Figure 5d**). Dominant taxa, including Actinobacteria and Proteobacteria (**Figure 5e**), tended to show similar relative abundance patterns in relic and total DNA over time, suggesting mortality but not decimation of these dominant microbes. Significant decreases in actinobacterial relative abundances between dry and wet soils in total DNA tracked with their patterns in relic DNA for the Jepson mound (but not swale) soils, and significant increases in proteobacterial relative abundances in response to wet-up were shared in relic and total DNA pools for both soils. These results are generally consistent with evidence for recent production of viruses of these dominant microbes throughout the experiment, and they suggest viral lysis as a possible mechanism for the production of new relic DNA both in dry (or drying) soils and during rewetting.

## Discussion

Here we show that the rewetting of seasonally dried soils – a critical event in Mediterranean grasslands that reactivates dormant soil microorganisms, leading to pulses of carbon and nitrogen mineralization^24^ – is consistently accompanied by a bloom of viral diversity, followed by extensive viral community turnover. Despite minimal overlap in viral community composition among the four soil types investigated in this study, successional dynamics were strikingly similar across soils. Substantial increases in both viral biomass and richness were observed within 24 hours of wet-up, with the magnitude increase in richness potentially dependent upon the length of the preceding dry period, suggesting that legacy effects play an important role in viral dynamics, consistent with prior evidence for historical effects on microbial community assembly^47^.

Viruses predicted to infect Actinobacteria and Proteobacteria (consistent key responders to the drying and rewetting of soils^25, 48^) largely followed the same trajectories as their hosts, suggesting that viruses prey on but do not necessarily decimate dominant microbial taxa in soil. The relative enrichment of actinobacteria under dry conditions has been reproducibly observed across multiple habitats^47, 49, 50^, including Mediterranean grasslands^25^, and here both actinobacteria and their putative phages were enriched in dry soils. Consistent with viral predation of actinobacteria (known drought-resistant microorganisms that can be active as soil dries^25, 48^) in dry soils, we recently reported an increase in putative actinophages upon decreasing soil moisture in a rainfall exclusion experiment^16^. In contrast, multiple Proteobacteria species have been identified as rapid responders to soil rewetting, often displaying the most active growth among microbial taxa during the first days following wet-up^30–34^. This expected increase in Proteobacteria was observed after rewetting here, together with a spike in putative proteobacteriophages. These results suggest that viruses respond to microbial activity upon rewetting, with substantial predation of the most abundant taxa. However, at least over the temporal and taxonomic scales considered here, no obvious ‘Kill-the-Winner’ dynamics (virus-driven decimation of dominant hosts^51^) were observed. Instead, host abundances remained relatively high when their viruses were abundant, with evidence of mortality of these abundant host taxa in the necromass pool. We suggest that soil viruses might ‘Cull-the-Winner’, preying on members of dominant, active populations but not entirely driving their hosts (and thereby themselves) to extinction.

Despite the general agreement in taxonomic trends between viruses and predicted hosts, viral communities displayed significantly higher rates of temporal succession than their prokaryotic counterparts. These differences are consistent with previously observed patterns in spatial structuring, whereby viral communities exhibited much more significant distance-decay relationships than prokaryotic communities^15, 16^. Biological and/or physicochemical explanations for the disconnect (*e.g.*, differences in phylogenetic scales of measurement^8, 52, 53^, an amplified signal in the viral communities due to large burst sizes, and/or differences in transport and attachment^9^) bear further exploration in future studies. However, we posit that these seemingly decoupled turnover patterns are also due in part to mismatched temporal windows of measurement for viruses in viromes and their hosts in total DNA. The extensive viral turnover here and the known short residence times for viral particles^54–58^ suggest that viromes capture a fleeting signal that reflects very recent activity. Measurements of soil microbial communities from total DNA, on the other hand, are known to include dormant organisms and relic DNA^37–39^, which can easily have residence times of months or longer^59^, such that a total DNA metagenome likely represents a much longer integrated period of time than its paired virome. Although these limitations of total DNA-based measurements are generally accepted and have been suggested to make minimal contributions to diversity estimates in some^37^ or most^39^ cases, the disconnected successional patterns observed here suggest that we might be missing more than previously appreciated, particularly during windows of very low and/or pulses of very high microbial activity (in dry and newly rewetted soils, respectively). Comparisons of active (e.g., RNA) and total (DNA) microbiomes^60^ are worth more extensive exploration over spatiotemporal scales in soil.

Substantial viral community diversity and heterogeneity over time, space, and environmental conditions are emerging as the rules in soil^11–18^, begging the question, how is this extreme viral diversity maintained? Viral survival in soils that experience seasonal drought, such as those studied here, is particularly mystifying. The assumption has been that viral populations piggyback on microbial dormancy (*e.g.*, vegetative states^61–63^) via lysogeny (mostly, viral genome integration into the host genome^64^)^5, 65–69^, but evidence is building against lysogeny as the primary viral survival mechanism in dry soils. Most known phages are exclusively lytic (virulent) and incapable of lysogeny, which requires specific genes for integration and excision from the host genome^64^. Thus far, lysogeny genes seem to be at low abundance in soil viromes^18, 70–72^, including in this study. However, results here also suggest that viral particle (virion) survival in dry soils is relatively rare (for example, two of our soils had dry soil virion abundances below detection limits), such that sustaining the extreme bloom of viral diversity upon wet-up from pre-existing virions alone would seem to require numerous successful infections sparked by few individuals spread across a highly diverse soil virosphere. Pseudolysogeny (arrested infection) may provide a viable alternative, allowing viral genomes to remain inside host cells, stalled in mid-infection. Although the name evokes lysogeny, pseudolysogeny is a state that can be achieved by virulent and temperate phages alike (i.e., phages that are exclusively lytic or capable of lysogeny)^73^. Stalled pseudolysogenic infections can resume when conditions allow, and they are a hallmark of phage infections under nutrient-limiting conditions in culture, including in soil *Bacillus* phage-host systems^74^. Pseudolysogeny thus offers the potential for lytic viral dormancy inside host cells, and it bears further study as a mechanism to enable viral persistence and seed banks that promote diversity in soil. Regardless of the dry season survival mechanism, the extreme viral turnover and production observed here - from viruses that largely do not possess recognizable lysogeny genes - suggests that exclusively lytic viruses may dominate in soil, potentially driving substantial microbial turnover, just as they do in the oceans^1, 75^.

Here, comparisons of DNase-treated and untreated viromes and total metagenomes have provided greater insights into viral and microbial dynamics and mortality than would be possible through any one approach alone. Despite some differences, the three sets of results (viral and prokaryotic composition in viromic and total DNA, respectively, and prokaryotic mortality patterns in relic DNA) generally converged, with relative abundances of hosts in both total and relic DNA and their putative viruses in viromes exhibiting similar patterns in response to wet-up. Both relic DNA abundance and the richness of the necromass pool decreased substantially within 24 hours of wet-up, indicating uptake and/or degradation of DNA from the dry season. Relic DNA re-accumulated over 10 days, indicating new microbial mortality, and both relic and total DNA composition tended towards convergence over time, suggesting increased similarity in the active and necromass pools in the days after wet-up. The mortality of microbes that continued to remain dominant in total DNA (including after an initial reduction in relic DNA removed most of the pre-existing dry soil necromass to reveal new mortality) is consistent with previous evidence for similarity in relic and total DNA diversity under some soil conditions^39^ and suggests both growth and death of these dominant taxa here. Thus, rather than the ‘Kill-the-Winner’ dynamics that might be expected to accompany extensive turnover in viral communities, these results are potentially consistent with a proposed ‘Cull-the-Winner’ model of viral predation that would cull dominant host taxa but not drive them to extinction. The extent to which this model might apply at different taxonomic, spatial, and eco-evolutionary scales requires further study and, particularly, improved virus-host linkages and a better understanding of host ranges in soil.

## Methods

### Soil collection

Soils used in our laboratory experiments were sourced from three grassland field sites in the Mediterranean climate zone of northern California: the University of California Hopland Research and Extension Center (Hopland, 39°00′14.6″N 123°05′09.1′W), the Donald and Sylvia McLaughlin Natural Reserve (McLaughlin, 38°52′32.1″N 122°25′09.7″W), and the Jepson Prairie Preserve (Jepson, 38°16′02.6″N 121°49′35.2″W). Since the terrain at the Jepson site is comprised of mima mounds and shallow swales that are edaphically distinct and support different plant communities, we collected separate soils from these two topographical features. For the main experiment, surface soils (depth: 0-15 cm) were harvested towards the end of the 2020 dry period, before the first seasonal rainfall: Hopland soils on October 29th, McLaughlin soils on October 30th, and Jepson soils on November 5th; for the preliminary experiment, surface soils from the Hopland site were harvested on November 16th, towards the end of the 2019 dry period. For Hopland and McLaughlin, we collected soils from four different locations, approximately 1 m apart from each other; for Jepson, we collected soils from four different mounds and, separately, their four adjacent swales. For each of the four soil types, we homogenized and sieved (8 mm) the four harvested soil cores and pooled them together into a composite soil sample, which was stored at room temperature in the dark until the soil microcosms were prepared.

### Laboratory simulations of wet-up

The main laboratory experiment was carried out in two separate batches: the first one included both Jepson soils and was started on November 24th, 2020; while the second one included the Hopland and McLaughlin soils and was started on January 6th, 2020. Microcosms were assembled by adding 10 g of dry soil to 50 mL bio-reaction centrifuge tubes with vented caps with 0.22 µm membranes inset in the caps (CELLTREAT #229476), a setup that enabled aseptic conditions while allowing free gas exchange. Molecular grade water was used to rewet dry soils to an average of 25% gravimetric soil moisture. Microcosms were then closed and kept at room temperature in the dark until destructively sampling them at 0 (pre-wet-up), 24, 48, 72, 120, 168, and 240 hours post-wet-up. To account for any changes under dry conditions during the 10-day time series, sets of non-wetted soil microcosms were harvested at the end of the experiment (T6 - dry). For each soil type, we set up replicated microcosms: 6 replicates for time points with paired DNase-treated and untreated viromes and 3 replicates for time points with only DNase-treated viromes (**Figure 1c**).

The preliminary experiment was performed on Hopland soils collected in the 2019 dry season and was started on February 12th, 2020. Three microcosms were assembled by adding 100 g of dry soil to 2.5” round pots (McConkey #JMCA25RSG) placed inside 1 L micropropagation containers with vented lids (Microbox #O119/140). Rewetting was performed by adding enough molecular grade water until soils were fully saturated. Microcosms were kept at room temperature in the dark until harvesting. Samples were collected from the same 3 microcosms at two time points: immediately before rewetting (pre-wet-up) and 336 hours post-wet-up.

### Total DNA extraction

Total DNA extractions were only performed for the main experiment. Immediately upon sample collection, we homogenized microcosms with a sterilized metal spatula and harvested a volume of soil approximately equivalent to the volume occupied by 250 mg of dry soil. For time points with paired DNase-treated and untreated viromes (**Figure 1c**), half volumes were separately collected for each microcosm and then pooled together. For both experiments, DNA was then extracted following the standard DNeasy PowerSoil Pro kit (Qiagen) protocol, except an additional 10-min 65 °C incubation was performed prior to bead-beating.

### Virome DNA extraction

Virome DNA extractions were performed on fresh soil immediately upon sample collection following a previously described protocol^16, 76^. For the main experiment, we added 10 mL of protein-supplemented phosphate-buffered saline solution (PPBS) directly into each microcosm tube immediately after collecting the soil volume used for total DNA extractions (see section above). For the preliminary experiment, we scooped two 10 g subsamples of soil from each microcosm (one subsample for DNase-treated viromes, the other one for untreated viromes) into 50 mL centrifuge tubes and added 10 mL of PPBS. Upon resuspending the soils via vortexing, the remaining steps were exactly the same for both experiments. Briefly, we mixed (10 min, 400 rpm, 4 °C in an orbital shaker) and centrifuged (10 min, 3,095 x g, 4 °C) the soil slurries and recovered the eluted virions suspended in the supernatant fraction. Soil pellets were back-extracted twice more, each time with 10 mL of fresh PPBS, and the resulting three ∼10 mL supernatants were pooled together. Soil particles remaining in these pooled supernatants were pelleted down via three rounds of centrifugation (10 min, 10,000 x g, 4°C), and the recovered supernatants were passed through a 0.22-µm polyethersulfone filter (MilliporeSigma) to deplete most cellular microorganisms. Recovered virions were concentrated by ultracentrifugation of the filtrates (2 h 25 min, 35,000 rpm, 4°C) and resuspension of the pellets in 100 µL of molecular grade water. For time points with paired DNase-treated and untreated viromes (**Figure 1c**), two 100-µL virion suspensions were mixed together at this stage before re-aliquoting into two separate 100-µL volumes to ensure that the resulting paired profiles were generated from the same pool of recovered virions. Untreated viromes were subjected directly to DNA extraction at this stage. For DNase-treated viromes, we incubated the resuspended virions with 10 units of RQ1 RNase-free DNase (Promega) for 30 minutes at 37 °C, then halted the reaction with 10 µL of stop solution. DNA extractions of both DNase-treated and untreated virion suspensions were performed with the DNeasy PowerSoil Pro kit (Qiagen) following the manufacturer’s protocol, except an additional 10-min 65°C incubation was performed prior to bead-beating. We used the Qubit dsDNA HS Assay to quantify the extracted DNA.

### Soil moisture measurement and soil chemistry profiling

For the main experiment, an additional set of six dry and six rewetted microcosms was set up for each soil type to track changes in soil moisture during the experiment. At each time point, we weighed the same set of microcosms and calculated soil moisture as the ratio of mass of water per mass of dry soil. Soil chemistry profiles of dry soils were generated by Ward Laboratories (Kearney, Nebraska, USA) and included the following measurements: soil pH and soluble salts (1:1 soil:water suspension), soil organic matter (percentage weight loss on ignition), nitrate (KCl extraction), potassium, calcium, magnesium, and sodium (ammonium acetate extraction), zinc, iron, manganese, and copper (DTPA extraction), phosphorus (Olsen method), and sulfate (Mehlich-3 extraction).

### Precipitation records

Total precipitation records from Jepson and McLaughlin were retrieved from the corresponding weather stations maintained by the Western Regional Climate Center (https://wrcc.dri.edu/). Records from Hopland were retrieved from the Sanel Valley weather station maintained by the Mendocino County (https://hrec.ucanr.edu/Weather,_Physical,_and_Biological_Data/).

### Shotgun library preparation, sequencing, and read quality-filtering

For both metagenomes and viromes, shotgun metagenomic libraries were constructed with the DNA Hyper Prep kit (Kapa Biosystems-Roche) and sequenced (paired-end, 150 bp) on the NovaSeq S4 platform (Illumina) by the DNA Technologies and Expression Analysis Core at the University of California, Davis Genome Center. Requested sequencing depth was approximately 20 Gbp per total metagenome and 10 Gbp per virome (both virome types), although the resulting depth varied drastically across virome profiles (**Supplementary Figure 12**). Reads were quality-filtered with Trimmomatic v0.33^77^, using a minimum q-score of 30 and a minimum read length of 50 bases. BBDuk v38.82^78^ was then used to remove any PhiX sequences.

### Amplicon library preparation, sequencing, and read quality-filtering

For the main experiment, we performed 16S rRNA gene amplicon sequencing on all total DNA samples following a previously described dual-indexing strategy^79, 80^. Briefly, we used the Platinum Hot Start PCR Master Mix (Thermo Fisher) and the 515F/806R set of universal primers to amplify the V4 region of the 16S rRNA gene. PCRs were performed following the Earth Microbiome Project’s^81^ PCR protocol: initial denaturation at 94 °C for 3 min; 35 cycles of 94 °C for 45 s, 50 °C for 60 s, and 72 °C for 90 s; and a final extension at 72 °C for 10 min. To identify any reagent contaminants, we generated a blank library by amplifying a control DNA template derived from a DNeasy PowerSoil Pro kit (Qiagen) extraction on molecular-grade water. Libraries were sequenced on the MiSeq platform (Illumina) by the DNA Technologies and Expression Analysis Core at the University of California, Davis Genome Center. To quality-filter reads, we used the filterAndTrim() function implemented in DADA2 v1.12.1^82^ with the following parameters: truncLen=c(0,0), maxN=0, maxEE=c(2,2), truncQ=2, rm.phix=TRUE.

### Identification and quantification of viral operational taxonomic units (vOTUs)

Sequence processing for viromes was performed separately for the main and preliminary experiments using the following workflow. MEGAHIT v1.2.9^83^ in metalarge mode was used to perform *de novo* assemblies of individual total metagenomes and viromes. For each assembly, we recovered all contigs with a minimum size of 10,000 bp and classifed them as viral with VIBRANT v1.2.1^84^, using the -virome flag for viromes but not for total metagenomes. High and medium quality viral contigs were then consolidated into species-level viral operational taxonomic units (vOTUs) with dRep v3.2.2^85^, using the single-linkage algorithm for hierarchical clustering and filtered nucmer alignments (≥95% ANI and ≥85% covered fraction, based on established benchmarks^41, 86^) for secondary clustering. We used Bowtie 2 v2.4.2^87^ in sensitive mode to map reads against the database of dereplicated vOTUs. Given the drastic differences in sequencing depth between some samples in the main experiment, (**Supplementary Figure 12**), we used the the view function implemented in SAMtools v1.11^88^ to subsample the resulting alignments. Briefly, we identified the lowest sequencing depth among post-wet-up libraries and divided it by the sequencing depth of each library to calculate a rarefaction factor for each alignment. Subsampling was performed independently for total metagenomes and viromes. For 2 pre-wet-up libraries with lower sequencing depth than the threshold used, no subsampling was performed. CoverM v.0.5.0 (https://github.com/wwood/CoverM) was used to parse subsampled alignments, quantify vOTU abundance and generate two coverage tables: one with the absolute number of aligned reads and the other one with the trimmed mean coverage. For both coverage tables, we used a ≥75% horizontal (breadth) coverage threshold to establish vOTU detection, as recommended by previously established benchmarks^86^. Finally, we discarded 726 vOTUs that were exclusively identified in single profiles (singletons).

### Gene-sharing network construction

We predicted the protein content of each vOTU with Prodigal v2.6.3^89^ (metagenome mode) and used the resulting amino acid file to build a gene-sharing network with vConTACT2 v0.9.19^42^ using Diamond^90^ to perform the protein alignment step and the MCL algorithm^91^ to identify protein clusters. The identified set of vOTU pairs was used to calculate a Fruchterman-Reingold network visualization layout with GGally v2.1.2^92^.

### Host taxonomy and temperate lifestyle predictions

Host predictions for vOTUs were originally attempted via a CRISPR-spacer-based approach. Briefly, we used Crass v1.01^93^ to assemble the CRISPR repeat and spacer arrays found in each sequenced library. We then used the BLASTn-short function implemented in BLAST v2.7.1^94^ to find spacer-protospacer matches against the dereplicated vOTU database (≤1 mismatch, e value < 1.0 x 10-10, and ≥95% identity) and repeat matches against the dereplicated MAG database (e value < 1.0 x 10-10 and 100% identity). Given the poor recovery of virus-host CRISPR-based linkages (**Supplementary Table 3**), we opted to use iPHoP v1.1.0^43^ with default parameters to predict the host taxonomy of the dereplicated vOTUs. Based on the gene annotations generated by VIBRANT v1.2.1^84^, we classified vOTUs as potential temperate viruses if there was at least one integrase or excisionase in their predicted protein content.

### Identification and quantification of metagenome-assembled genomes (MAGs)

The quality of the *de novo* assemblies generated from individual total metagenomes (see section above) was low (on average, we recovered 300 >10 Kbp contigs per assembly). As such, we opted to follow a co-assembly approach to facilitate the binning process. We used MEGAHIT v1.2.9^83^ in metalarge mode (with a minimum contig length of 2 Kbp) to perform eight co-assemblies, each one including either all of the total metagenomes or all of the DNase-treated viromes of a specific soil type. Read recruitment against each co-assembly was performed with Bowtie 2 v2.4.2^87^ in sensitive mode, and contig coverage depth was quantified with the jgi_summarize_bam_contig_depths function implemented in MetaBAT 2^95^. Contigs were then binned into metagenome assembled genomes (MAGs) with VAMB v3.0.2^96^, and MAG quality was assessed with CheckM v1.1.3^97^. We recovered all ≥ 50 Kbp MAGs with ≥ 50% completeness and ≤ 10% contamination and de-replicated them with dRep v3.2.2^85^, using the single-linkage algorithm for hierarchical clustering and filtered nucmer alignments (default settings of ≥99% ANI and ≥10% covered fraction) for secondary clustering. We used Bowtie 2 v2.4.2^87^ in sensitive mode to map reads against the database of dereplicated MAGs and subsampled the resulting alignments using the same strategy described above for vOTUs. We then used CoverM v.0.5.0 (https://github.com/wwood/CoverM) to quantify MAG abundance and generate two coverage tables: one with the absolute number of aligned reads and the other with the trimmed mean coverage. For both coverage tables, we used a ≥25% horizontal (breadth) coverage threshold to establish MAG detection. Finally, we discarded 2 MAGs that were exclusively identified in single profiles.

### Taxonomic classification and phylogenetic tree construction

We used the classify_wf pipeline implemented in GTDB-Tk v2.1.0^98^ to identify bacterial and archaeal marker genes in the dereplicated set of MAGs, perform multiple sequence alignments (MSA), and assign taxonomic classifications. The resulting MSA file was further used to build a phylogenetic tree via the GTDB-Tk infer workflow using default parameters.

### 16S rRNA gene amplicon sequence processing

We used DADA2 v1.12.1^82^ (default settings) to perform error rate inference, dereplication, denoising, read merging, and chimera removal. Taxonomy assignments were performed with the DADA2 implementation of the RDP classifier^99^, using the GTDB database v207^100^ as a reference. Upon discarding any amplicon sequence variant (ASV) detected in the blank library, we rarified the libraries to the minimum sequencing depth observed across samples (23,054 reads). Finally, we discarded 19,456 ASVs that were exclusively identified in single profiles.

### Statistical analyses

All statistical analyses were performed with R v3.6.3^101^ with heavy reliance on the tidyverse v1.3.2^102^ suite of packages for data wrangling. Unless otherwise noted, analyses were performed on trimmed mean coverage tables. Where applicable, multiple comparison corrections were performed with the Bonferroni method. We used vegan v2.5-7^103^ to calculate Bray-Curtis dissimilarities on log-transformed relative abundances and Jaccard indexes on presence/absence tables. We performed principal coordinates analyses with ape v5.4.1^104^ and set intersection analyses with eulerr v6.1.1^105^ and UpSetR v1.4.0^106^. Temporal distance-decay relationships were calculated via Pearson’s correlation tests and linear regressions evaluating the association between temporal distance and Bray-Curtis similarity across pairs of samples. To identify vOTUs and MAGs with differential abundance patterns, we used DESeq2 v1.38.1^107^ to run negative binomial generalized models testing the effect of time point (coded as a discrete variable) on taxon abundance. For vOTUs, differential abundance analyses were exclusively performed on DNase-treated profiles. For all DESeq2 analyses, we used non-normalized count tables as input and discarded all taxa with extreme count outliers via DESeq2’s default Cook’s distance threshold. Temporal trend groups were defined via hierarchical clustering of standardized abundances (z-score) using the average linkage algorithm. Enrichment analyses in local network neighborhoods were performed following an implementation of the Spatial Analysis of Functional Enrichment (SAFE) algorithm^108^ that was previously used to assess trait distributions in vOTU gene-sharing networks^16^. Briefly, we used igraph v1.3.5^109^ to determine the local neighborhood of each node in the network by identifying all nodes that could be reached (either directly or indirectly) via a path with a length shorter than the first percentile of the collection of shortest paths connecting all possible pairs of nodes in the network. To assess the overrepresentation or underrepresentation of temporal groups in each local neighborhood, we performed hypergeometric tests using the phyper() function with the “lower.tail” parameter set to false and true, respectively. Local neighborhoods smaller than 10 nodes were discarded before correcting for multiple comparisons. Hypergeometric tests were also used to evaluate the overrepresentation and underrepresentation of microbial phyla in each vOTU and MAG trend group, discarding all phyla with less than 10 members in the dataset. We used emmeans v1.8.3^110^ to perform Dunnett’s tests assessing differences in the aggregated abundances of Actinobacteriota and Proteobacteria MAGs in post-wetup time points relative to dry soils. Plots were generated with ggplot2 v3.4.0^111^, ggtree v3.6.2^112^, eulerr v6.1.1^105^, and UpSetR v1.4.0^106^.

## Data availability

Raw sequences are available at the NCBI Sequence Read Archive (BioProject PRJNA859194). The databases of dereplicated vOTUs and MAGs are available at 10.5281/zenodo.7510627. Code and processed files needed to replicate the statistical analyses can be found at https://github.com/cmsantosm/WetupViromes.

## Acknowledgements

This work was predominantly supported by an award from the U.S. Department of Energy, Office of Science, Office of Biological and Environmental Research, Genomic Science Program, #DE-SC0020163 (grant to JPR and JBE). Additional support was provided by the U.S. Department of Energy, Office of Science, Office of Biological and Environmental Research, Genomic Science Program, award #DE-SC0021198 (grant to JBE). Research at Lawrence Livermore National Laboratory was conducted under the auspices of DOE Contract DE-AC52-07NA27344. Shotgun metagenomic library construction and high-throughput sequencing were performed by the DNA Technologies and Expression Analysis Core at the UC Davis Genome Center, supported by NIH Shared Instrumentation Grant 1S10OD010786-01. We thank the University of California Natural Reserves site directors and staff, including Shane Waddell, Cathy Koehler, Jeffrey Clary, and John Bailey, for facilitating access to field sites and associated site information.

## Supplementary Figures

**Supplementary Figure 1.**
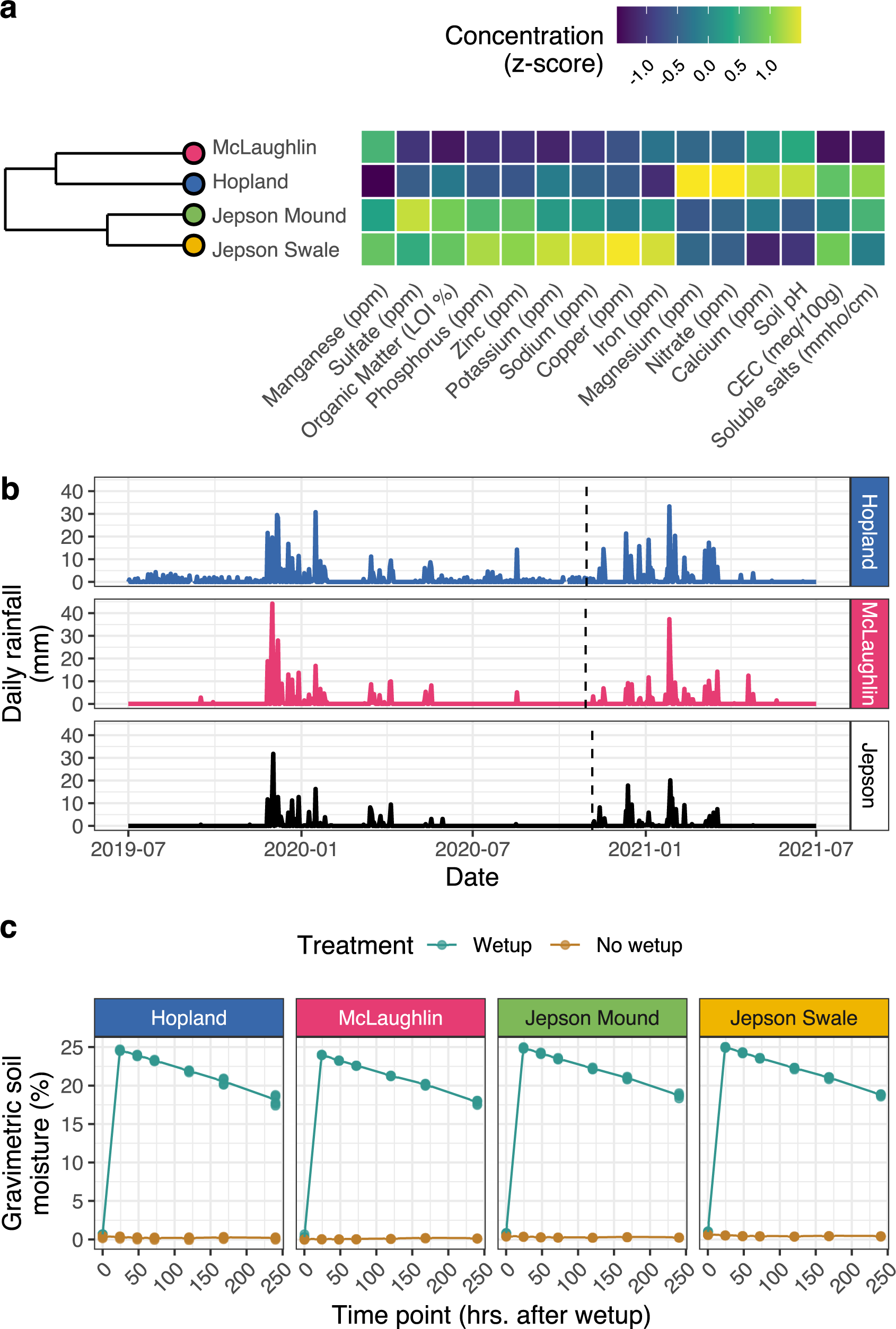
(a) Hierarchical clustering of soil types based on their abiotic properties. The heatmap shows the z-transformed values of the edaphic variables included in the analysis. **(b)** Daily precipitation at each of the three field sites preceding, during, and after soil harvesting. Dotted vertical lines mark the dates of soil sample collection. **(c)** Gravimetric soil moisture content over time across soil types. Points correspond to individual microcosms, and trend lines represent the mean moisture content at each time point. Color indicates whether replicates underwent rewetting or remained dry.

**Supplementary Figure 2.**
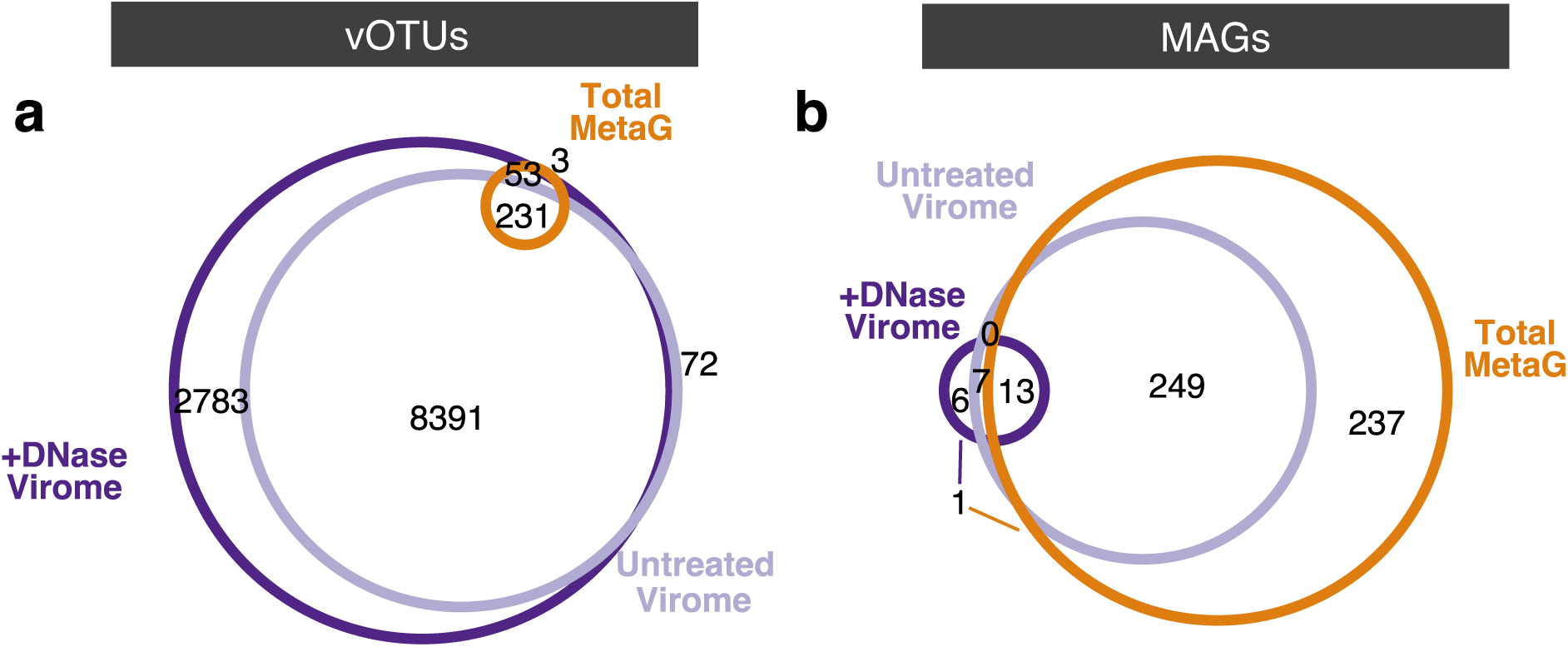
**(a-b)** Euler diagrams showing the intersections between the sets of **(a)** vOTUs and **(b)** MAGs detected in each profiling method.

**Supplementary Figure 3.**
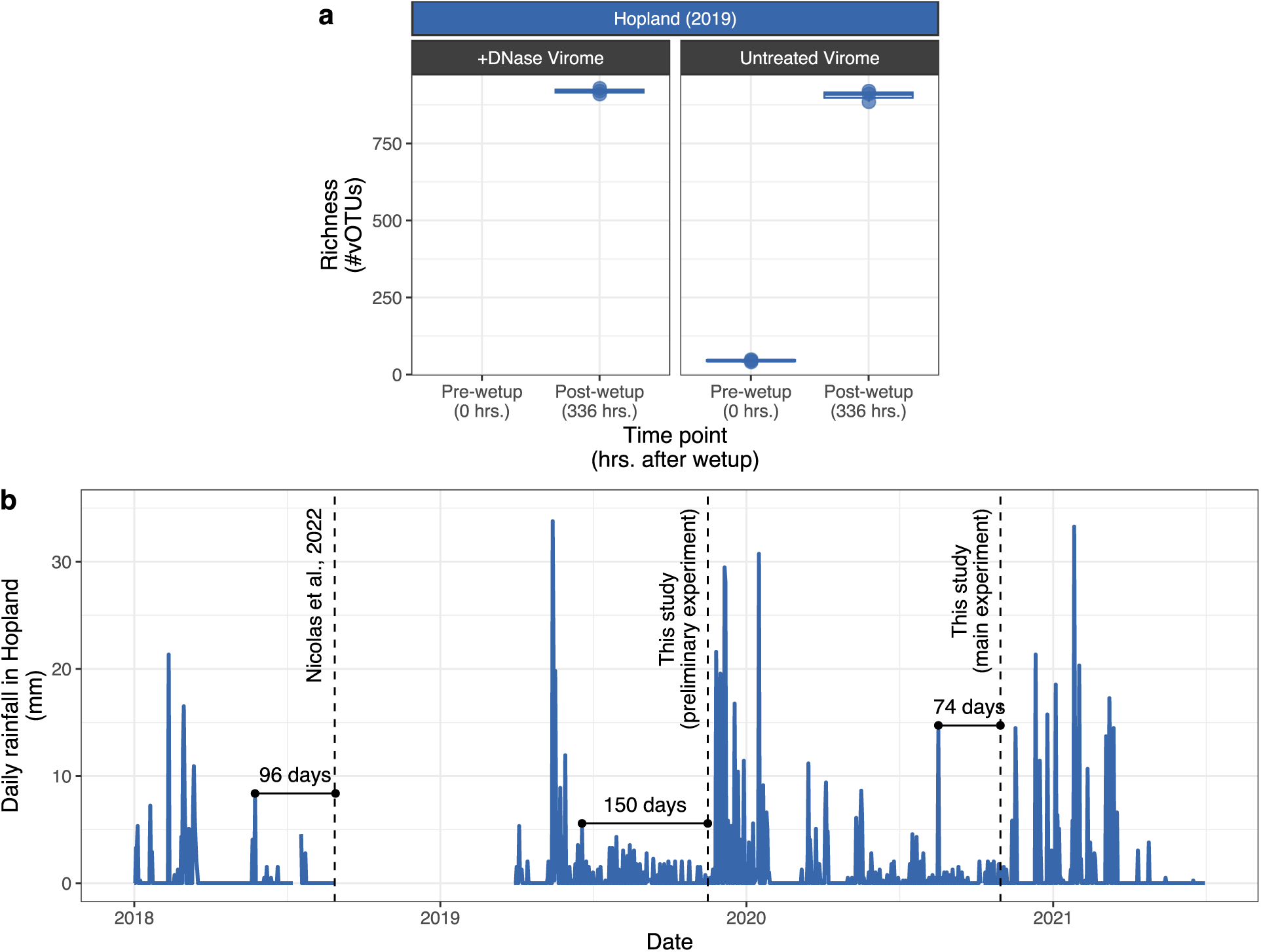
**(a)** Viral community richness in a preliminary viromic survey of dry and rewetted (336 hrs after a laboratory simulation of wet-up) soils harvested from the Hopland site at the end of the 2019 dry season (approximately one year prior to sample collection for the main microcosm experiment in this study). Facets correspond to DNase-treated and untreated viromic profiles, and boxes display the median and interquartile ranges. DNase-treated profiles from dry soils could not be generated due to low DNA yields. **(b)** Daily precipitation at the Hopland site between 2018 and 2021. Dotted vertical lines mark the dates of soil sample collection for the main (2020) and preliminary (2019) wet-up experiments reported in this study, as well as the wet-up experiment from 2018 reported in Nicolas et al., 2022. Black horizontal lines highlight the period since the most recent ≥ 5 mm rainfall event preceding each soil sample collection time point. Gaps in the trend line between 2018 and 2019 indicate missing data from the weather station.

**Supplementary Figure 4.**
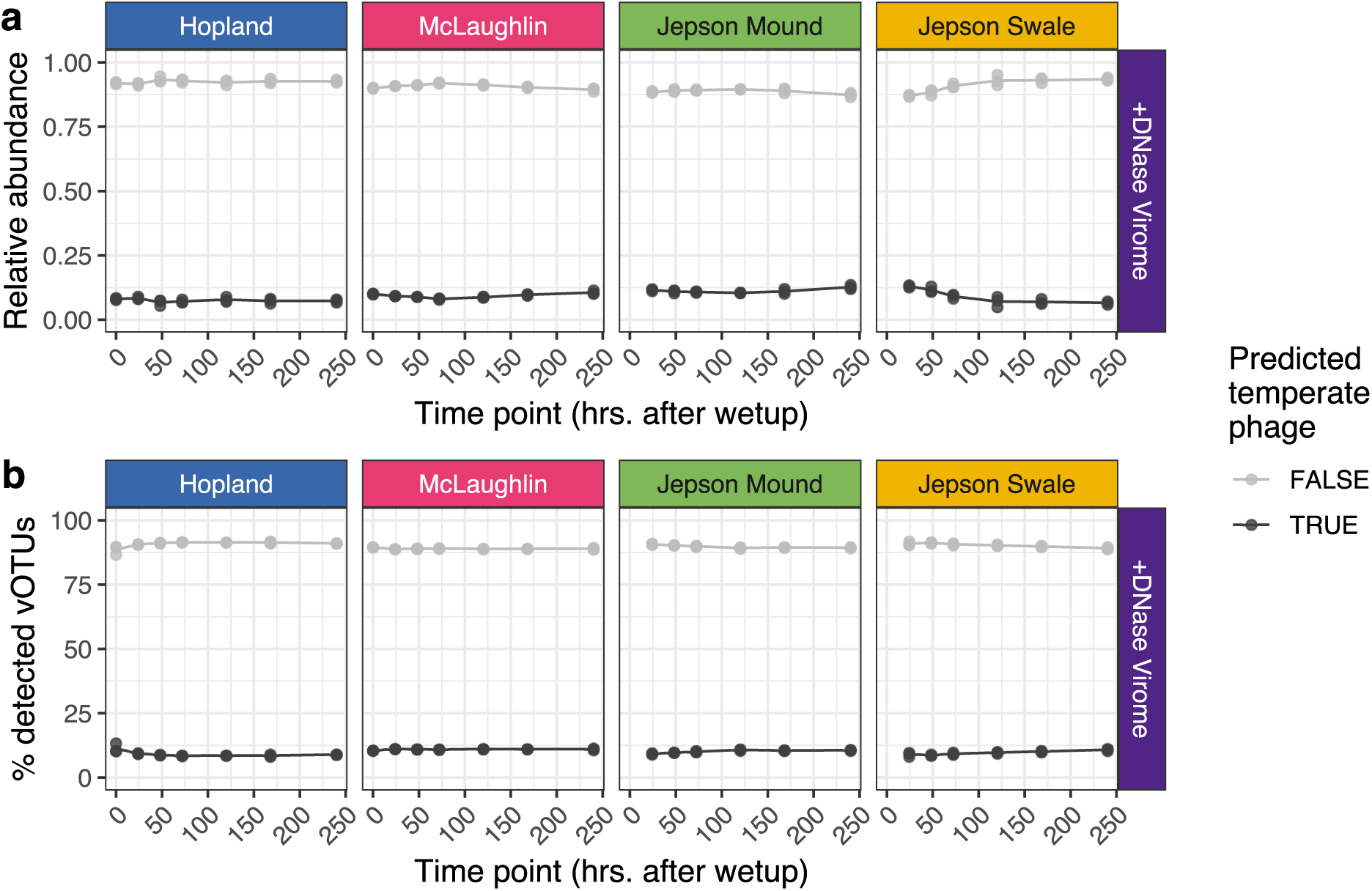
Proportions of predicted temperate phages (phages putatively capable of lysogeny) and phages not predicted to be temperate in viral communities by soil type. Trends for all vOTUs are separated according to their temperate phage prediction (true = predicted temperate phage, false = not predicted to be temperate). For each time point, panel **(a)** shows aggregated vOTU relative abundances (based on read mapping-derived coverage), and panel **(b)** shows the percentage of total vOTUs detected. Points are individual samples, and trend lines track the mean values among replicates.

**Supplementary Figure 5.**
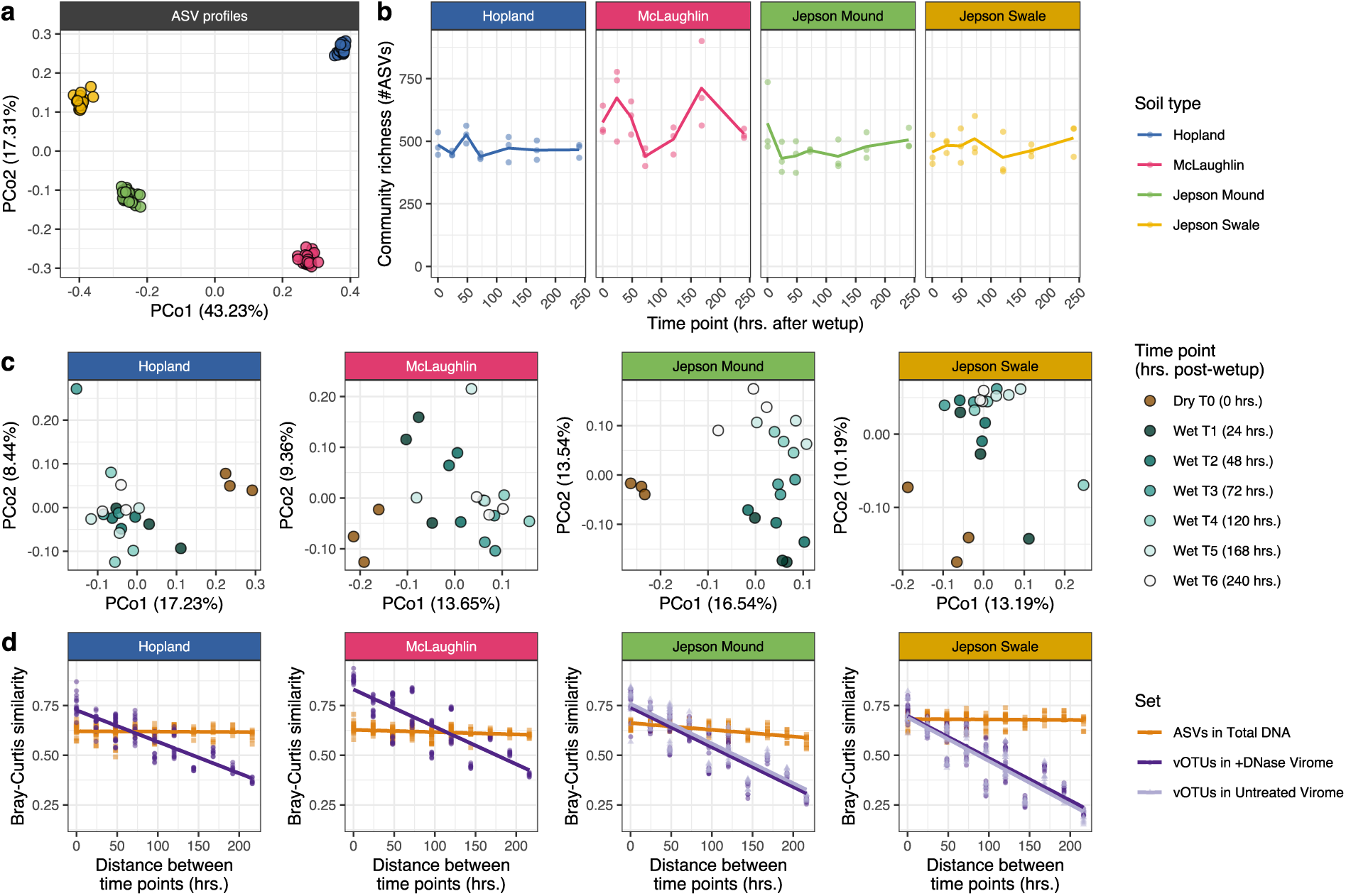
Compositional patterns of prokaryotic communities based on ASV profiles of the 16S rRNA gene from total DNA (a complement to trends for MAGs reported in main text figures). **(a)** Unconstrained analysis of principal coordinates performed on Bray-Curtis dissimilarities. Colors indicate soil type. **(b)** Community richness measured as the number of ASVs recovered from soil microcosms: colored points correspond to replicates and colored trend lines represent the mean richness at each collection time point. **(c)** Unconstrained analyses of principal coordinates performed ASV Bray-Curtis dissimilarities. Analyses were performed independently for each soil type (facets). Colors indicate the collection time point and soil status (dry or wet). **(d)** Linear regressions between Bray-Curtis similarities and temporal distances in post-wet-up time points for viral (vOTUs in viromes) and prokaryotic (ASVs in total DNA) communities. Points represent pairs of samples, trend lines show the least squares linear regression model, and facets correspond to soil types.

**Supplementary Figure 6.**
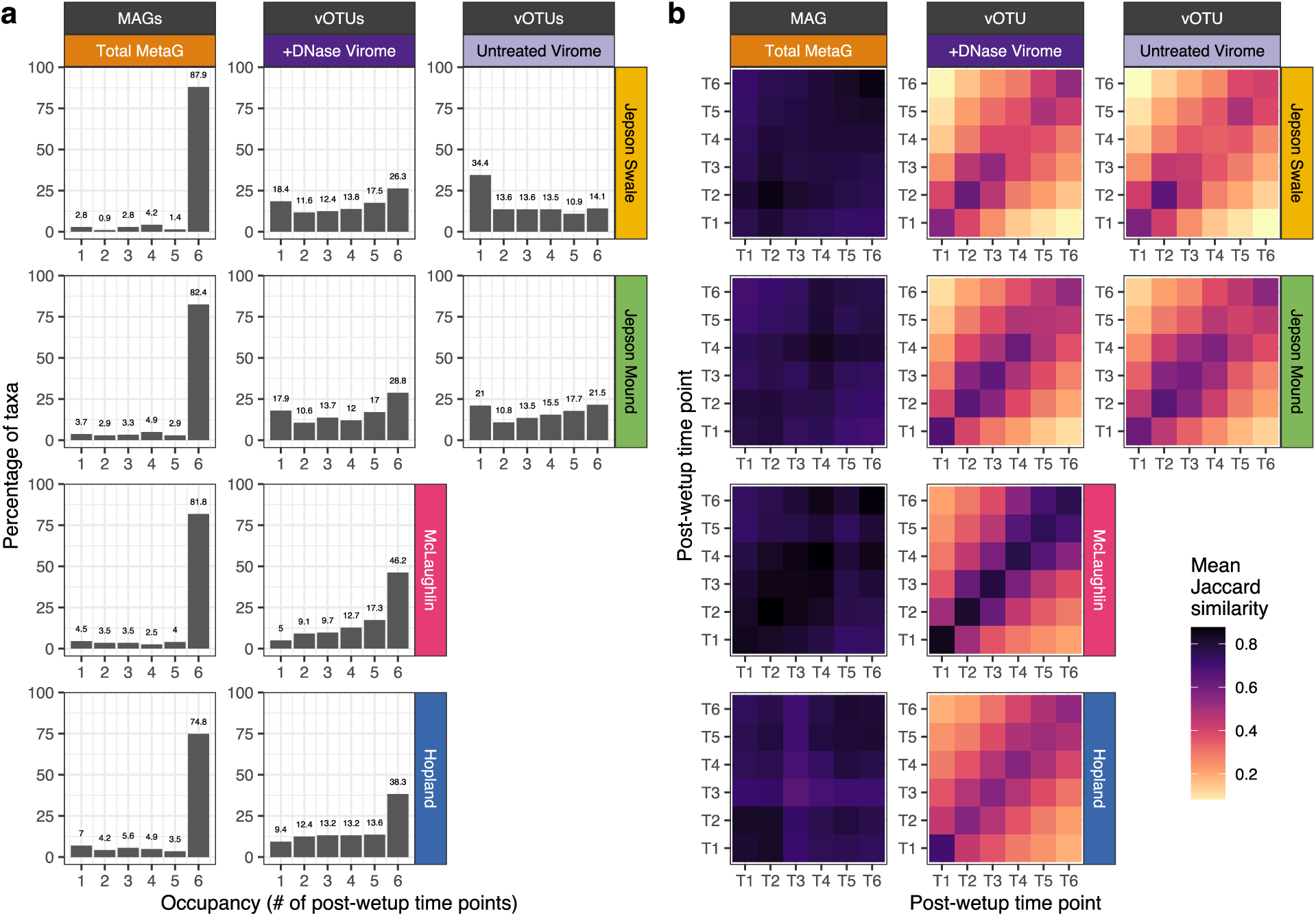
**(a)** Percentage of MAGs (left) and vOTUs (right) detected at each occupancy level (number of post-wet-up time points) in total metagenomes and viromes, respectively. **(b)** Pairwise Jaccard similarities of MAG communities profiled in total metagenomes (left) and vOTU communities profiled in DNase-treated and untreated viromes (right). The x-axis and y-axis are the same, such that the diagonal from lower left to upper right for each facet indicates pairwise community similarity among technical replicates, and other comparisons are between microcosms at different time points.

**Supplementary Figure 7.**
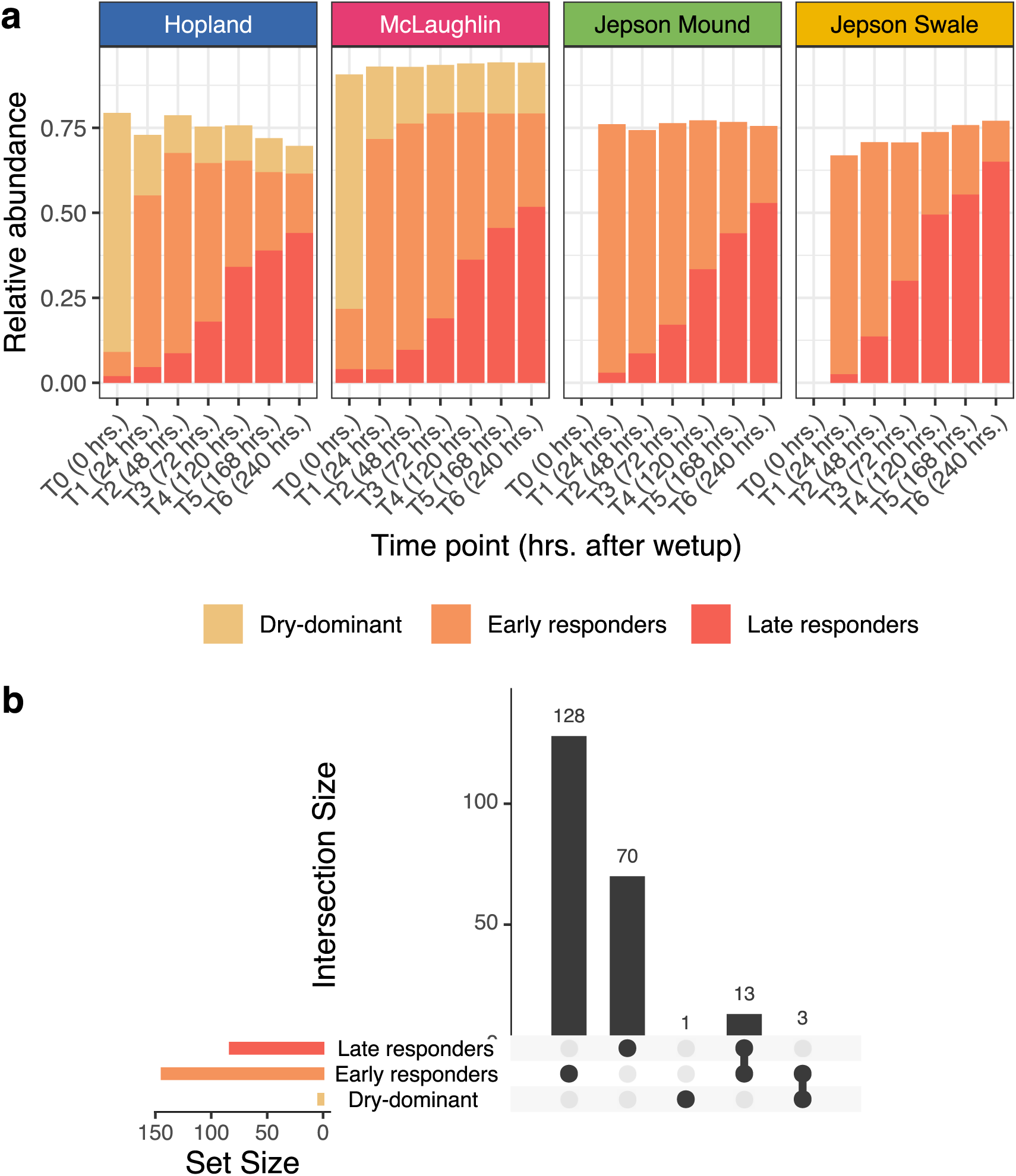
**(a)** Changes in the aggregated abundances of vOTUs in each trend group. **(b)** Upset plot displaying the temporal groups assigned to vOTUs detected as differentially abundant across multiple soils: light gray vertical bars correspond to vOTUs displaying the same temporal trend across soil types, while dark gray vertical bars correspond to vOTUs displaying different temporal trends across soil types.

**Supplementary Figure 8.**
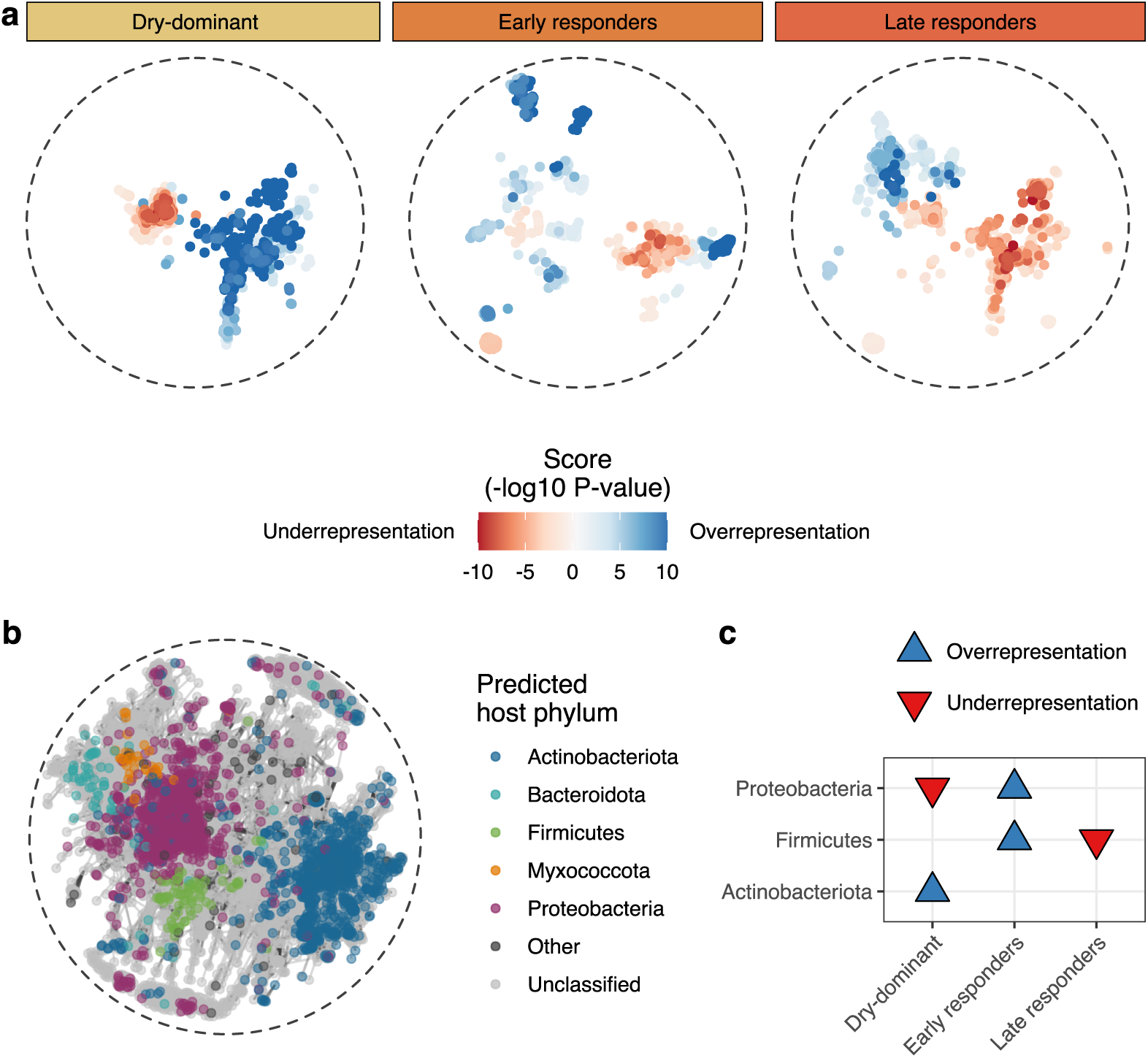
Viral taxonomic network analyses. For **(a-b)**, the network layout was constructed with the Fruchterman-Reingold algorithm, based on pairs of vOTUs (nodes) with a significant overlap in their predicted protein content. **(a)** Distribution of local network neighborhoods with a significant overrepresentation (blue points) or underrepresentation (red points) of vOTUs assigned to a particular trend group (adjusted p-value < 0.05, hypergeometric test). Each colored point denotes the center of a significant local neighborhood, and the color gradient indicates the extent of the significance of the overrepresentation or underrepresentation of the temporal group in that neighborhood. **(b)** Distribution of predicted hosts in the gene-sharing network. For each vOTU (node), the predicted host phylum is indicated. **(c)** Set of predicted host phyla significantly overrepresented or underrepresented in each vOTU temporal group (adjusted p-value < 0.05, hypergeometric test).

**Supplementary Figure 9.**
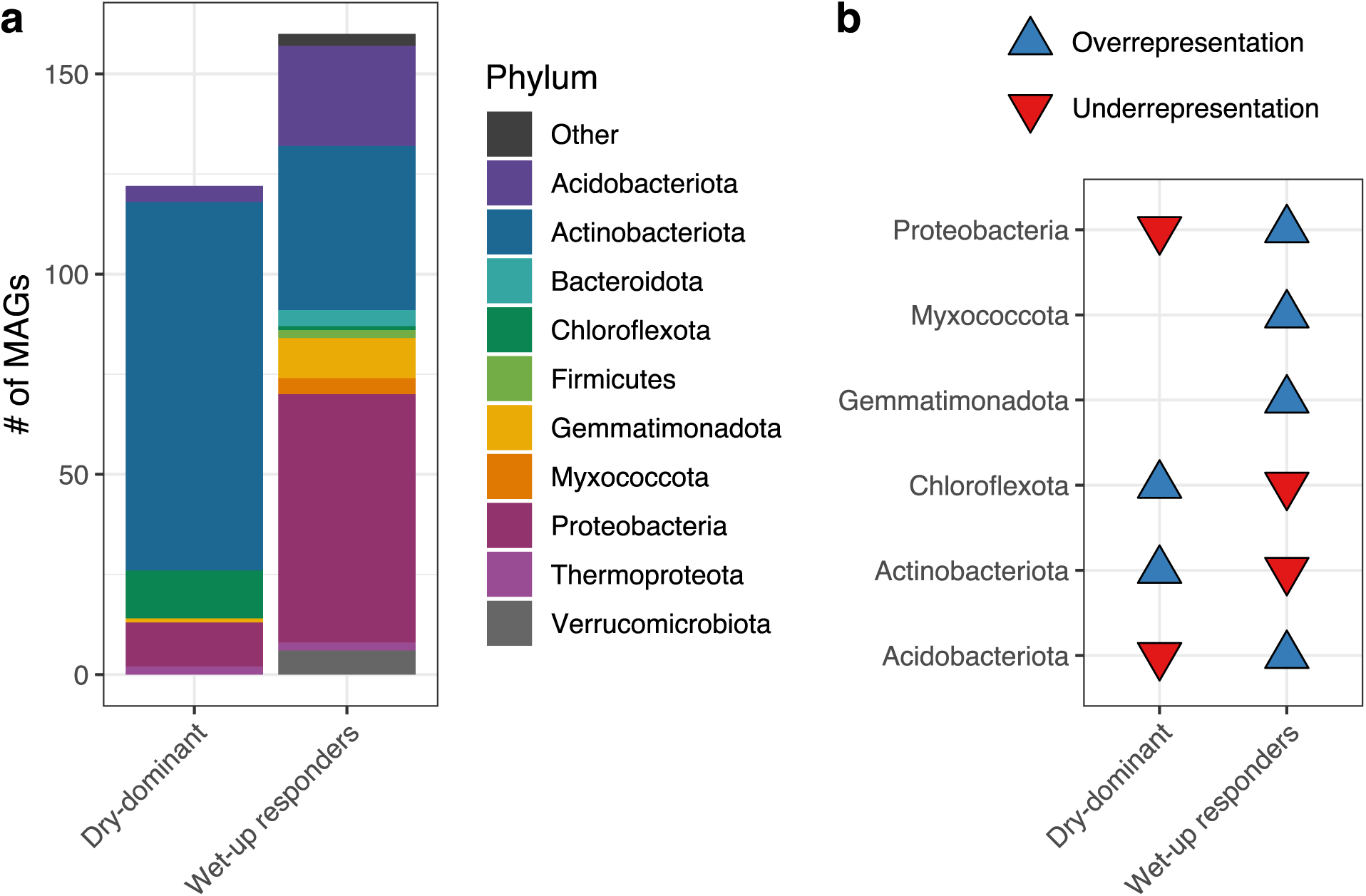
**(a)** Taxonomic composition of MAG temporal groups. **(b)** Set of MAG phyla significantly overrepresented or underrepresented in each MAG temporal group (adjusted p-value < 0.05, hypergeometric test).

**Supplementary Figure 10.**
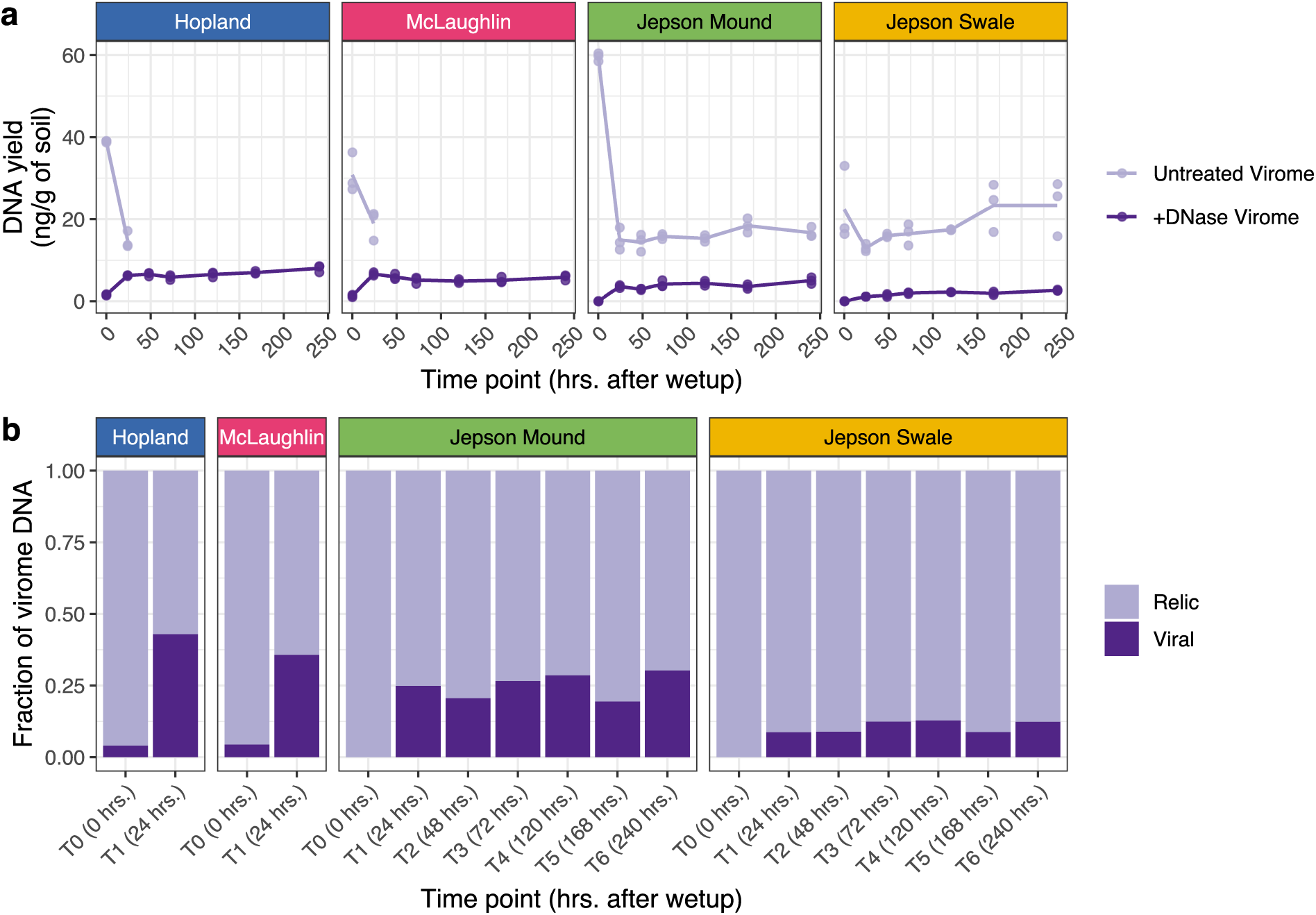
**(a)** Temporal trends in the DNA yields recovered from DNase-treated and untreated viromes extracted from soil microcosms. Points correspond to individual samples, and trend lines track mean yields among technical replicates over time. Yields were below detection limits for pre-wet-up samples from Jepson soils (represented as zero in the graph). **(b)** Mean proportion of relic DNA (*i.e.*, DNA that could be digested by a DNase treatment) and viral DNA in the viral-size-fraction recovered acrossn replicates at each time point. In panel **(b)**, the fraction of viral DNA was calculated by dividing the yield of each DNase-treated virome by the yield of its paired untreated virome.

**Supplementary Figure 11.**
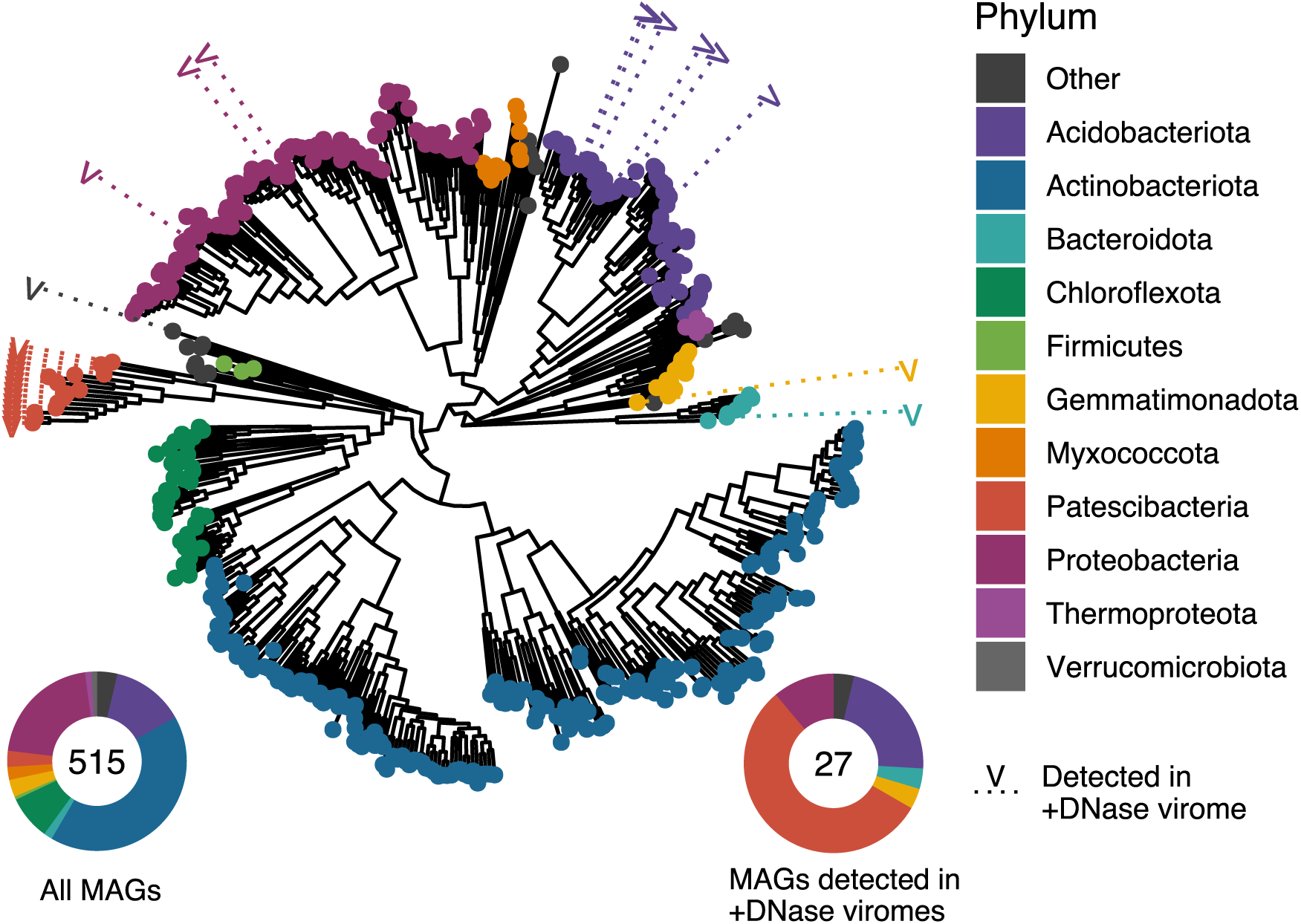
**(a)** Phylogenetic tree displaying the set of MAGs recovered across all samples. Tip color indicates MAG taxonomic classification at the phylum level. MAGs recovered in DNase-treated virome profiles are highlighted with a “V” label.

**Supplementary Figure 12.**
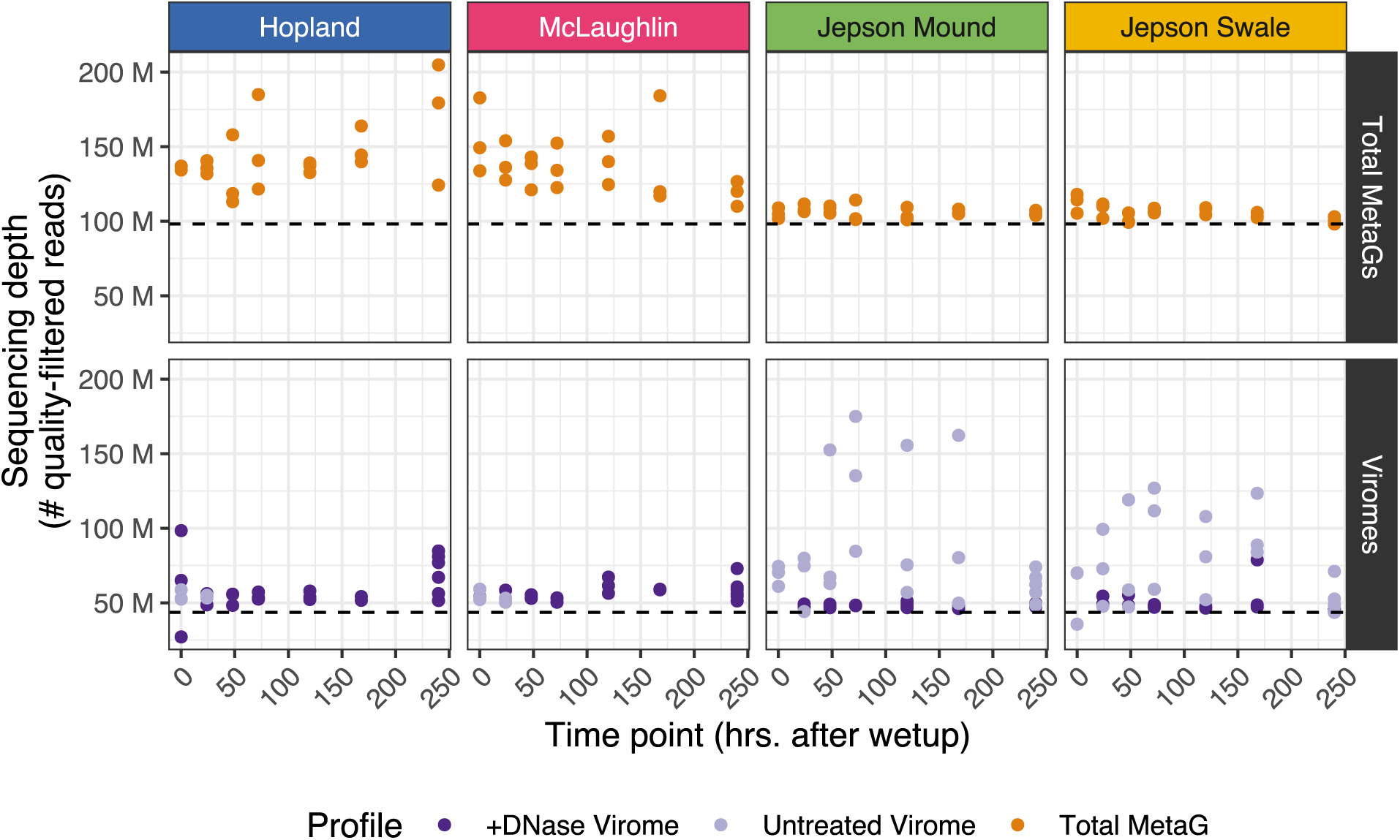
Sequencing depth (measured as number of reads) of quality-filtered libraries. Horizontal dotted lines indicate the sequencing depth used as a reference to calculate alignment rarefaction factors for total metagenomes (top facet) and viromes (bottom facet).

## Supplementary Tables

**Supplementary Table 1.** Permutational multivariate analyses of variance (PERMANOVA) testing differences in vOTU community composition between dry soil microcosms collected at T0 and dry soil microcosms collected at T6. Tests were performed on DNase-treated viromes for Hopland and McLaughlin soils and on untreated viromes for Jepson soils.

| Soil | Profile | Formula | Df | SumsOfSqs | MeanSqs | F.Model | R2 | Pr(>F) |
| --- | --- | --- | --- | --- | --- | --- | --- | --- |
| Jepson Mound | Untreated viromes | dist ~ Timepoint | 1 | 0.090 | 0.090 | 2.250 | 0.360 | 0.3 |
| Jepson Swale | Untreated viromes | dist ~ Timepoint | 1 | 0.105 | 0.105 | 2.012 | 0.335 | 0.1 |
| Hopland | DNase-treated viromes | dist ~ Timepoint | 1 | 0.025 | 0.025 | 2.133 | 0.348 | 0.1 |
| McLaughlin | DNase-treated viromes | dist ~ Timepoint | 1 | 0.010 | 0.010 | 5.559 | 0.582 | 0.1 |

**Supplementary Table 2.**
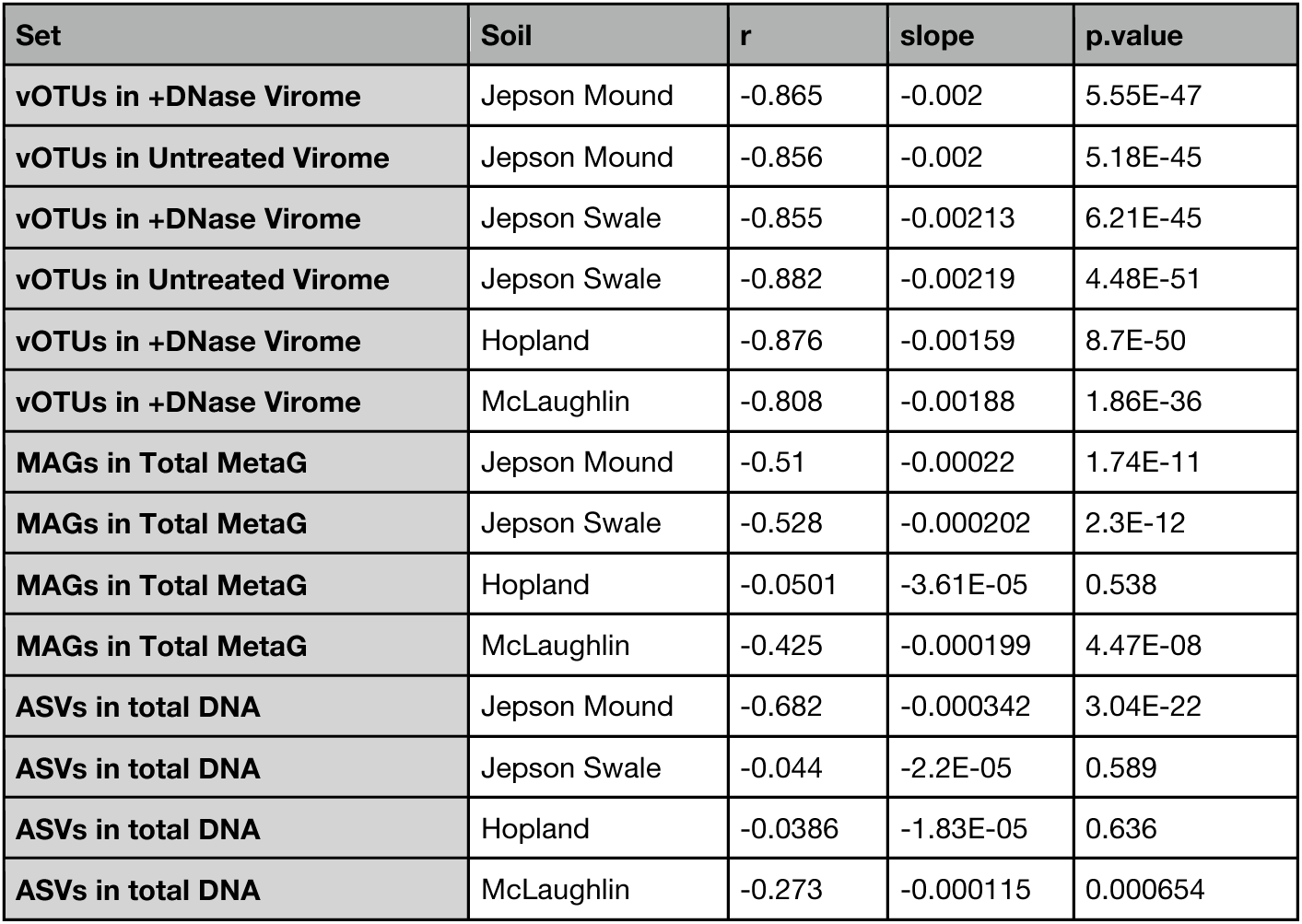
Relationship between vOTU/MAG/ASV Bray-Curtis similarity and temporal distance in post-wet-up time points for each soil type. Columns show the Pearson’s correlation coefficient (r), the linear regression slope, and the associated P-value. Analyses were performed independently for MAG communities profiled in total metagenomes, vOTU communities profiled in DNase-treated viromes, and vOTU communities profiled in untreated viromes.

**Supplementary Table 3.** List of predicted virus-host linkages found via CRISPR spacer/protospacer matches. A total of 22 vOTUs were linked to 5 MAGs. Taxonomic classification of MAGs is provided in the last 7 columns.

| vOTU_ID | MAG_ID | Domain | Phylum | Class | Order | Family | Genus | Species |
| --- | --- | --- | --- | --- | --- | --- | --- | --- |
| WUA_VD34_contig_126882 | JB_10374 | Bacteria | Chloroflexota | Ktedonobacteria | Ktedonobacterales | Ktedonobacteraceae | Unclassified | Unclassified |
| WUA_VN42_contig_192307 | JB_10374 | Bacteria | Chloroflexota | Ktedonobacteria | Ktedonobacterales | Ktedonobacteraceae | Unclassified | Unclassified |
| WUA_VN45_contig_76662 | JB_10374 | Bacteria | Chloroflexota | Ktedonobacteria | Ktedonobacterales | Ktedonobacteraceae | Unclassified | Unclassified |
| WUB_VD25_contig_61940 | JB_20884 | Bacteria | Proteobacteria | Alphaproteobacteria | Acetobacterales | Acetobacteraceae | Palsa-883 | Unclassified |
| WUA_VD35_contig_5163 | JB_5390 | Bacteria | Myxococcota | Myxococcia | Myxococcales | 40CM-4-68-19 | 40CM-4-68-19 | Unclassified |
| WUA_VD36_contig_24611 | JB_5390 | Bacteria | Myxococcota | Myxococcia | Myxococcales | 40CM-4-68-19 | 40CM-4-68-19 | Unclassified |
| WUA_VD37_contig_99257 | JB_5390 | Bacteria | Myxococcota | Myxococcia | Myxococcales | 40CM-4-68-19 | 40CM-4-68-19 | Unclassified |
| WUA_VD39_contig_85096 | JB_5390 | Bacteria | Myxococcota | Myxococcia | Myxococcales | 40CM-4-68-19 | 40CM-4-68-19 | Unclassified |
| WUA_VD40_contig_12344 | JB_5390 | Bacteria | Myxococcota | Myxococcia | Myxococcales | 40CM-4-68-19 | 40CM-4-68-19 | Unclassified |
| WUA_VN25_contig_170948 | JB_5390 | Bacteria | Myxococcota | Myxococcia | Myxococcales | 40CM-4-68-19 | 40CM-4-68-19 | Unclassified |
| WUA_VN32_contig_217547 | JB_5390 | Bacteria | Myxococcota | Myxococcia | Myxococcales | 40CM-4-68-19 | 40CM-4-68-19 | Unclassified |
| WUB_VD07_contig_22197 | JB_5390 | Bacteria | Myxococcota | Myxococcia | Myxococcales | 40CM-4-68-19 | 40CM-4-68-19 | Unclassified |
| WUB_VD13_contig_78199 | JB_5390 | Bacteria | Myxococcota | Myxococcia | Myxococcales | 40CM-4-68-19 | 40CM-4-68-19 | Unclassified |
| WUB_VD29_contig_53475 | JB_5390 | Bacteria | Myxococcota | Myxococcia | Myxococcales | 40CM-4-68-19 | 40CM-4-68-19 | Unclassified |
| WUB_VD35_contig_103068 | JB_5390 | Bacteria | Myxococcota | Myxococcia | Myxococcales | 40CM-4-68-19 | 40CM-4-68-19 | Unclassified |
| WUB_VD37_contig_2291 | JB_5390 | Bacteria | Myxococcota | Myxococcia | Myxococcales | 40CM-4-68-19 | 40CM-4-68-19 | Unclassified |
| WUB_VN11_contig_42049 | JB_5390 | Bacteria | Myxococcota | Myxococcia | Myxococcales | 40CM-4-68-19 | 40CM-4-68-19 | Unclassified |
| WUA_VN32_contig_104725 | JT_2215 | Bacteria | Acidobacteriota | Acidobacteriae | Acidobacteriales | Gp1-AA117 | Gp1-AA17 | Unclassified |
| WUA_VD41_contig_38616 | JT_6980 | Bacteria | Actinobacteriota | Actinomycetia | Mycobacteriales | Jatrophihabitantaceae | Iso899 | Unclassified |
| WUA_VN38_contig_226568 | JT_6980 | Bacteria | Actinobacteriota | Actinomycetia | Mycobacteriales | Jatrophihabitantaceae | Iso899 | Unclassified |
| WUB_VD10_contig_75231 | JT_6980 | Bacteria | Actinobacteriota | Actinomycetia | Mycobacteriales | Jatrophihabitantaceae | Iso899 | Unclassified |
| WUB_VD35_contig_69775 | JT_6980 | Bacteria | Actinobacteriota | Actinomycetia | Mycobacteriales | Jatrophihabitantaceae | Iso899 | Unclassified |

